# Damage-sensing recruitment of a lipid phosphatase couples lysosomal membrane repair to proteostatic adaptation

**DOI:** 10.64898/2026.04.04.716461

**Authors:** Yanwei Su, João Mello-Vieira, Dmytro Puchkov, Gillian Dornan, Max Ruwolt, Elisabeth Südhoff, Oluwatobi Andrew Adeosun, Heike Vogel, Simon Sündermann, Annette Schürmann, Joost C. M. Holthuis, Fan Liu, Ivan Dikic, Michael Ebner, Volker Haucke

## Abstract

Restoration of organellar membrane integrity is critical for maintaining cellular homeostasis. Lysosomal membrane damage activates local repair machineries and global stress responses, but how signaling lipid metabolism is engaged by damage sensors to support and mechanistically link these processes remains poorly understood. Here we show that the phosphoinositide 3-phosphatase MTMR14 is recruited to damaged lysosomes through calcium-dependent binding to sphingomyelin. At these sites, MTMR14 promotes local PI(3)P hydrolysis and supports PI(4)P accumulation, thereby facilitating formation of ER–lysosome contact sites associated with membrane repair, without affecting ESCRT recruitment. MTMR14-dependent lipid remodelling causes reduced mTORC1 signalling and a decrease in global protein synthesis, consistent with an acute proteostatic adaptation to lysosomal injury. Cells lacking MTMR14 display impaired damage-induced lipid remodelling, altered repair-associated structures, sustained protein synthesis, and increased sensitivity to lysosomal injury, all of which can be mitigated by mTORC1/S6K inhibition. Our findings identify damage-sensing recruitment of MTMR14 and local PI(3)P turnover on damaged lysosomes as a phosphoinositide module that promotes lysosomal membrane integrity and homeostasis while functionally linking nutrient signalling to proteostasis under membrane stress.

---

Cells and tissues experience membrane injury from mechanical and chemical stress, invading pathogens, and the accumulation of peptide or protein aggregates that can perforate organellar membranes. The ability to cope with such damage is essential for viability and organismal health ^1-4^. Defects in membrane damage responses have been implicated in human disease, including muscular dystrophies, neurodegeneration, cancer and cardiovascular disease ^5^. Lysosomes are particularly vulnerable targets because they concentrate hydrolases and redox-active metabolites, and they function as central hubs for catabolism, nutrient sensing and quality control ^6-8^.

Lysosomal membrane permeabilization (LMP) can be triggered by lysosomotropic agents, membrane-active peptides, aggregation-prone proteins such as neuropathogenic tau fibrils, or intracellular pathogens that exploit the endolysosomal system ^9-13^. Moreover, lysosomal membrane damage via release of iron can induce specific cell-death pathways such as ferroptosis^14^. Hence, lysosomal membrane damage is relevant to neurodegeneration, muscular dystrophy^5^, aging, and infection and can be a critical vulnerability of cancer cells^10, 12^.

To survive LMP, cells engage a multilayered lysosomal damage response comprising membrane remodelling, rapid repair, selective removal and regeneration of lysosomes ^4, 6, 10, 12, 15-17^. Limited damage elicits fast local repair processes, including endosomal sorting complex required for transport (ESCRT)-dependent sealing of endolysosomal membranes labelled by Galectins^18-20^ and lipid transfer–mediated repair via endoplasmic reticulum (ER)–lysosome contact sites, in which phosphatidylinositol 4-phosphate [PI(4)P] on damaged lysosomes recruits lipid transfer proteins to drive cholesterol and phospholipid delivery (the phosphoinositide-initiated membrane tethering and lipid transport, or PITT, pathway) ^21, 22^. Additional mechanisms include Ca^2+^-activated sphingomyelin scrambling and hydrolysis at lysosomes ^23^, conjugation of ATG8 proteins to single membranes (CASM) at damaged endolysosomes ^24, 25^, and formation of stress granules that can physically plug and stabilize lesions ^10, 26^. When damage is irreparable, endolysosomes are marked by ubiquitin ^27^ and targeted for selective autophagy (lysophagy) ^10, 12, 15, 28^, followed by lysosome biogenesis and membrane regeneration to restore degradative capacity.

These local damage responses intersect with broader signalling networks. Canonical mTORC1 signalling on lysosomes integrates nutrient and growth factor cues to control protein synthesis and autophagy ^4, 29, 30^, and is modulated by lysosomal phosphoinositides and organelle positioning ^31-33^. We and others have shown previously that phosphatidylinositol 3-phosphate [PI(3)P] on endolysosomal membranes promotes mechanistic target of Rapamycin (mTORC1) activity ^10, 12, 15, 28^, whereas repression of mTORC1 is required to initiate ULK1-dependent autophagy and lysophagy ^4, 12^. Non-canonical pathways, including transcription factor EB (TFEB)-mediated transcriptional programs, couple lysosomal Ca^2+^ release to lysosome biogenesis and autophagic lysosome reformation. In addition, Galectins act as damage sensors: Galectin-8 recruits the GALTOR complex to inhibit mTORC1 on damaged lysosomes ^34^, while Galectin-3 and Galectin-8 also participate in damage sensing ^35^ and lysophagy initiation, e.g. during infection ^36^. Protein translation control via mTORC1 and the lysosomal degradative machinery constitute a major cellular proteostatic network ^37-39^. How proteostasis is integrated with these distinct regulatory circuits—ESCRT and PITT-mediated repair, CASM and stress-granule–dependent stabilization, lysophagy, and mTORC1/TFEB/Galectin signalling—and how they are coordinated in space and time at damaged lysosomes remains incompletely understood.

Phosphoinositides are prime candidates to provide such spatiotemporal coordination. These low-abundance lipids define organelle identity and regulate membrane traffic, signalling and contact sites ^40, 41^. PI(3)P and PI(4)P are of particular relevance to the lysosomal damage response: lysophagy and autophagic clearance of damaged lysosomes require PI(3)P generated by VPS34, whereas PI(4)P is essential for PITT-mediated ER–lysosome contacts and repair ^21, 22^, and also contributes to lysophagy. ESCRT components typically bind PI(3)P-rich membranes ^42^, although ESCRT recruitment to damaged lysosomes can be maintained even when class III PI 3-kinase activity is inhibited ^18^, suggesting that distinct phosphoinositide pools or additional signals may operate at these sites. Recent studies reported that PI(4)P accumulates on damaged lysosomes ^21, 22, 43^. We have shown that, under nutrient stress, a phosphoinositide conversion pathway on lysosomes links PI(3)P to PI(4)P and mTORC1 signalling: MTMR14 (also known as Jumpy), a disease-associated PI(3)P 3-phosphatase ^44-46^, modestly accumulates on lysosomes in starved cells to attenuate canonical mTORC1–S6K signalling ^47^. Whether similar lipid circuits operate in response to lysosomal membrane damage, whether phosphoinositide phosphatases engage as part of the immediate damage response, and to what extent they contribute to repair, lysophagy and proteostasis is unknown.

Here we investigate how local lipid signaling shapes the lysosomal response to membrane damage. We identify damage-sensing recruitment of MTMR14 and local PI(3)P turnover on damaged lysosomes as a local phosphoinositide module that promotes lysosomal membrane integrity and function while functionally linking nutrient signalling to proteostasis under membrane stress.

## Membrane damage induces rapid loss of lysosomal PI(3)P via the PI 3-phosphatase MTMR14

The various pathways for lysosomal repair and removal of damaged membranes are known to exhibit differential signaling lipid requirements (Figure 1a): Membrane repair by lipid transfer via the PITT pathway depends on lysosomal PI(4)P ^21, 22^, while lysophagy requires PI(3)P, a lipid shown to stimulate nutrient signaling via mTORC1^31, 47^, which in turn is known to repress lysophagy ^30^. ESCRT proteins typically associate with organellar membranes enriched in PI(3)P^42^, although in the case of lysosomal membrane damage ESCRT recruitment appears to be unaffected by PI 3-kinase inhibition^18^.

**Figure 1.**
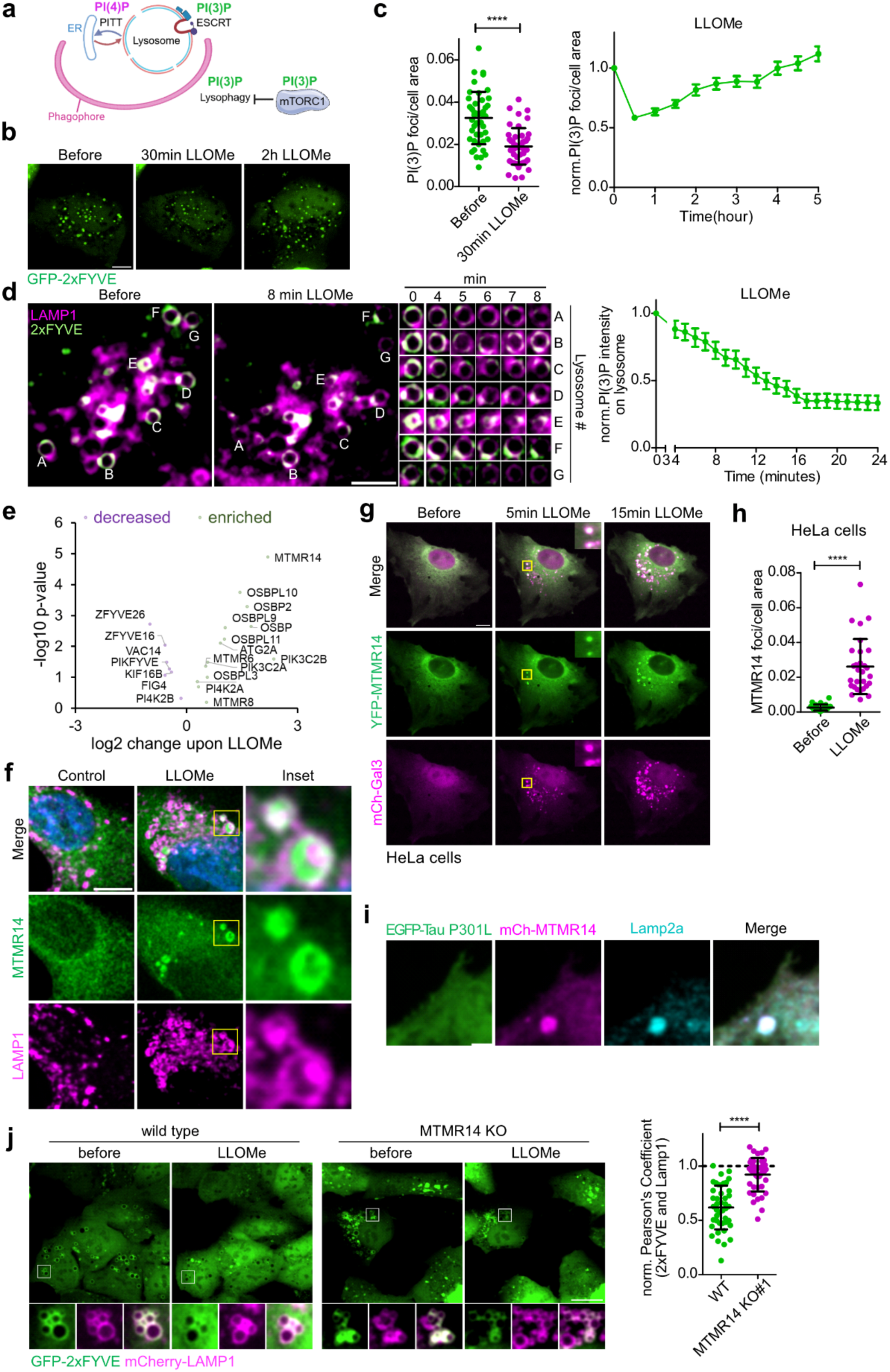
LMP recruits MTMR14 to regulate PI(3)P levels on damaged lysosomes. (a) Schematic illustration of putative roles of PI(4)P and PI(3)P in lysosome repair and removal pathways. ESCRT, endosomal sorting complex required for transport. PITT, phosphoinositide-initiated membrane tethering and lipid transport pathway. ER, endoplasmic reticulum. mTORC1, mechanistic target of rapamycin complex 1. (b) Biphasic PI(3)P dynamics in 0.5mM LLOMe-treated cells. The same HeLa cell expressing eGFP-2xFYVE was imaged before and after different durations of LLOMe treatment. Scale bar, 10 µm. (c) Left: quantification of (b), number of PI(3)P puncta/cell area before and after 30 minutes of 0.5mM LLOMe was compared by t test (n = 39 cells). Right: time course of PI(3)P change in 0.5mM LLOMe treated HeLa cells expressing eGFP-2xFYVE. Quantification of the number of PI(3)P puncta/cell area, fold change over control. (n = 39 cells). Data are mean ± s.e.m. (d) LLOMe decreases PI(3)P on lysosomes. Left: magnification of the perinuclear region of a U2OS cell co-expressing eGFP-2xFYVE and mCherry-LAMP1 imaged after different durations of 0.5mM LLOMe treatment. Capital letters indicate same vesicles before and after 8 minutes of LLOMe treatment. Scale bar, 5 µm. Middle: time-resolved changes of GFP-2xFYVE on the same mCherry-LAMP1 vesicles as indicated in the left panels by capital letters. Right: Representative time course (mean±s.e.m.) of LMP-induced PI(3)P decline on lysosomes from ten cells. See also Fig. 1j and Extended Data Fig. 1b for quantifications. (e) Volcano plot illustrating the enrichment of phosphoinositide kinases, phosphatases, phosphoinositide-binding proteins, and lipid-transport proteins in Lyso-IP samples from LLOMe-treated cells. Lysosomes were immunoprecipitated from control or 30 minutes 1mM LLOMe-treated HEK293 cells expressing TMEM192-3xHA and subjected to quantitative MS/MS analysis. (f) Representative confocal images of mScarlet-MTMR14 CRISPR-Cas9 knockin U2OS cells treated with control (DMSO) or 15 minutes 1mM LLOMe. Cells were co-stained with antibodies specific for mScarlet (MTMR14) and LAMP1. Scale bar, 10 µm. (g) Representative confocal images of HeLa cells co-expressing YFP-MTMR14 with mCherry-Galectin3 imaged live before and after 5 and 15 minutes of 1mM LLOMe treatment. Scale bar, 10 µm. (h) Quantification of the number of MTMR14 puncta/cell from control and 1 h 1mM LLOMe treated HeLa cells. t test (n = 29 cells). (i) Expression of EGFP-Tau P301L recruits MTMR14 to lysosomes. Shown are magnifications of confocal images of HeLa cells co-expressing EGFP-Tau P301L with mCherry-MTMR14 and stained with antibodies against Lamp2a. Scale bar, 2 µm. (j) Left: representative confocal live cell images of wild type and MTMR14 KO U2OS cells co-expressing GFP-2xFYVE with mCherry-LAMP1 imaged before and after 20 minutes of 0.5mM LLOMe treatment. Right: quantification of the Pearson’s Coefficient of GFP-2xFYVE and mCherry-LAMP1 from LLOMe treated U2OS WT and MTMR14 KO cells before and after 20 minutes LLOMe treatment. Dotted line denotes Pearson’s Coefficient of GFP-2xFYVE and mCherry-LAMP1 before LLOMe set to 1. Scale bar, 20 µm. t test, n=43 cells in U2OS WT and 41 cells in U2OS MTMR14 KO cells. Statistical analyses were performed using GraphPad Prism. Two-tailed unpaired t-test, paired t-test or one-sample t-tests were conducted using column statistics to compare the sample means to a hypothetical value of 1 or one-way ANOVA with Tukey’s multiple comparisons test. All bar graphs represent mean ± SD unless otherwise stated. ***p < 0.001, **p < 0.01, *p < 0.05. See also Extended Data Fig. 1 and 2.

We hypothesized that PI(3)P, which marks endosomes as well as signaling-active lysosomes in fed cells at steady-state ^47^, may be subject to membrane damage-induced regulation. To test this, we first monitored the levels and dynamics of endolysosomal PI(3)P in HeLa cells expressing eGFP-2xFYVE as a probe^47, 48^ by sensitive Yokogawa W1 live spinning disk confocal microscopy. Induction of lysosomal membrane damage (hereafter referred to as lysosomal membrane permeabilization, LMP) by the lysosomotropic agent l-leucyl-l-leucine methyl ester (LLOMe) triggered a moderate overall reduction of PI(3)P puncta (Figure 1b), followed by gradual recovery of PI(3)P over the next hours (Figure 1b,c). LLOMe-induced membrane damage requires cathepsin activity and is thus restricted to degradative lysosomes^49^. We therefore focused our further analysis on PI(3)P on lysosomes marked by mCherry-LAMP1 and found that LMP triggered a rapid (i.e. within 3-5 min) and pronounced reduction of lysosomal PI(3)P reaching steady-state low PI(3)P levels within 15 min (Figure 1d). Comparison of LLOMe treatment and pharmacological PI(3)P kinase inhibition using VPS34-IN1 suggest that LLOMe leads to an approximately 50% reduction of lysosomal PI(3)P (Extended Data Figure 1b). Cytoplasmic eGFP-2xFYVE intensity mildly increased or remained unchanged during the imaging period of 24 min (Extended Data Fig. 1a), suggesting that the apparent loss of lysosomal PI(3)P was not caused by photobleaching of the eGFP-2xFYVE probe (as further confirmed genetically, see Fig. 1j, below). Lysosomal PI(3)P reduction was mirrored by a concomitant rapid and sustained rise in lysosomal PI(4)P levels monitored by eGFP-OSBP-PH (Extended Data Fig. 1c), in agreement with previous data ^21, 22^. The levels of phosphatidylinositol 3,5-bisphosphate [PI(3,5)P_2_], a signaling lipid specific to the endolysosomal system, were largely unaffected upon damage (Extended Data Fig. 1d), although small effects of this lipid on the cellular response to LMP cannot be ruled out (as suggested by recent data ^50^). Lysosomal membrane damage, thus, triggers a rapid reduction of PI(3)P at lysosomes.

To unravel the underlying mechanism, we followed an unbiased proteomic approach by lysosome-immunoprecipitation from HEK293 cells expressing TMEM192-3xHA with or without LLOMe treatment followed by quantitative mass spectrometry analysis (Table S1). Several members of the oxysterol-binding protein family (OSBPs) as well as ATG2A (i.e. components of the PITT pathway) previously reported to be recruited to damaged lysosomes ^21,22^ were highly enriched on lysosomes from LLOMe-treated cells (Figure 1e). In addition, we observed a differential enrichment of phosphoinositide kinases (e.g. phosphatidylinositol 4-kinase 2α, PI4K2A) and phosphatases on damaged lysosomes (Figure 1e). The most enriched of these enzymes was the phosphoinositide 3-phosphatase MTMR14, an enzyme implicated in diseases including myopathy, spermatogenesis^45, 51, 52^, and cancer^46, 53-55^ (Figure 1e). Other members of the MTMR family of lipid phosphatases were either not (i.e., MTM1) or only mildly enriched (i.e., MTMR8, MTMR6) on lysosomes from LLOMe-treated cells (Extended Data Figure 1e,f). Damage-induced lysosomal recruitment of native endogenous mScarlet-MTMR14^endo^ was confirmed by confocal microscopy in genome-engineered knock-in U2OS cells stained with mScarlet-specific antibodies (Figure 1f). Further analysis by live spinning disk confocal imaging demonstrated that membrane damage triggered the rapid recruitment (i.e. within 5 min) of YFP-MTMR14 to permeabilized lysosomes marked by mCherry-Galectin3 in HeLa cells (Figure 1g, h; Movie S1), U2OS osteosarcoma cells (Extended Data Figure 1g), in C2C12 myoblasts (Extended Data Figure 1h), or in microglial BV2 and HMC3 cells (Extended Data Figure 1I,j). Damage-induced MTMR14 recruitment to lysosomes was dose-dependent and could be triggered by concentrations as low as 100 µM LLOMe (Extended Data Figure 1k).

These data suggest that MTMR14 is an early and sensitive responder to LMP. Other MTMR family members were recruited much less effectively (Extended Data Figure 1f). Alternative inducers of LMP such as the lysosomotropic detergent MSDH or the peptide glycyl-L-phenylalanine 2-naphthylamide (GPN) also led to MTMR14 recruitment to lysosomes (Extended Data Figure 2a,b). Furthermore, MTMR14 recruitment to lysosomes was induced by osmotic or oxidative stress but not by mitochondrial uncoupling, alkalinization of the endolysosomal system in presence of Bafilomycin A1, or the H^+^/K^+^-ionophore Nigericin (Extended Data Figure 2c). Interestingly, permeabilization of the plasma membrane by benzalkonium chloride (BAC) triggered MTMR14 recruitment to the cell surface (Extended Data Figure 2c). Finally, we probed the effects of mimicry of neuropathogenic tau protein aggregates that occur in Alzheimer’s disease by overexpressing mutant EGFP-tau (P301L). Expression of mutant tau (P301L) led to a time-dependent accumulation of damaged lysosomes marked by Galectin3 (Extended Data Figure 2d) that colocalized with co-expressed MTMR14 puncta at LAMP2a-positive lysosomes in U2OS cells (Figure 1i).

To test a possible functional role of MTMR14 in PI(3)P depletion at damaged lysosomes, we genome-engineered MTMR14 knockout (KO) U2OS cells and verified successful KO by newly generated MTMR14-specific antibodies (Extended Data Figure 2e). The damage-induced hydrolysis of global PI(3)P as well as PI(3)P on lysosomes observed in wild-type (WT) cells was nearly completely abrogated in KO cells lacking MTMR14 (Figure 1j, Extended Data Figure 2f,g).

Hence, membrane damage induces the rapid loss of lysosomal PI(3)P mediated by recruitment of the PI 3-phosphatase MTMR14.

## Rapid lysosomal membrane recruitment of MTMR14 via calcium-triggered association with sphingomyelin

Next, we aimed to dissect the mechanism that enables MTMR14 recruitment to damaged lysosomes. First, we subjected MTMR14 to deletion mutagenesis. This analysis identified a C-terminal fragment of MTMR14 that was required for recruitment to lysosomes in LLOMe-treated cells (Extended Data Figure 3a). Alpha Fold 2 prediction suggests that this C-terminal fragment of MTMR14 comprises an α-helix containing clusters of hydrophobic residues that may interact with the hydrophobic core of the damaged lipid bilayer embedded into an intrinsically disordered region. Indeed, when fused to mCherry the C-terminal fragment of MTMR14 was sufficient for targeting to damaged lysosomes where it colocalized with full length YFP-MTMR14 (Figure 2a). Further kinetic analysis revealed the rapid recruitment of the mCherry-fused MTMR14 C-terminal domain with similarly fast kinetics (i.e. within 3 min) as PI(4)P monitored by eGFP-OSBP-PH as a sensor (Figure 2b; Movie S2). Of note, both probes largely overlapped on the same organelles, suggesting that lysosomal PI(3)P loss and local PI(4)P synthesis may be mechanistically linked.

**Figure 2.**
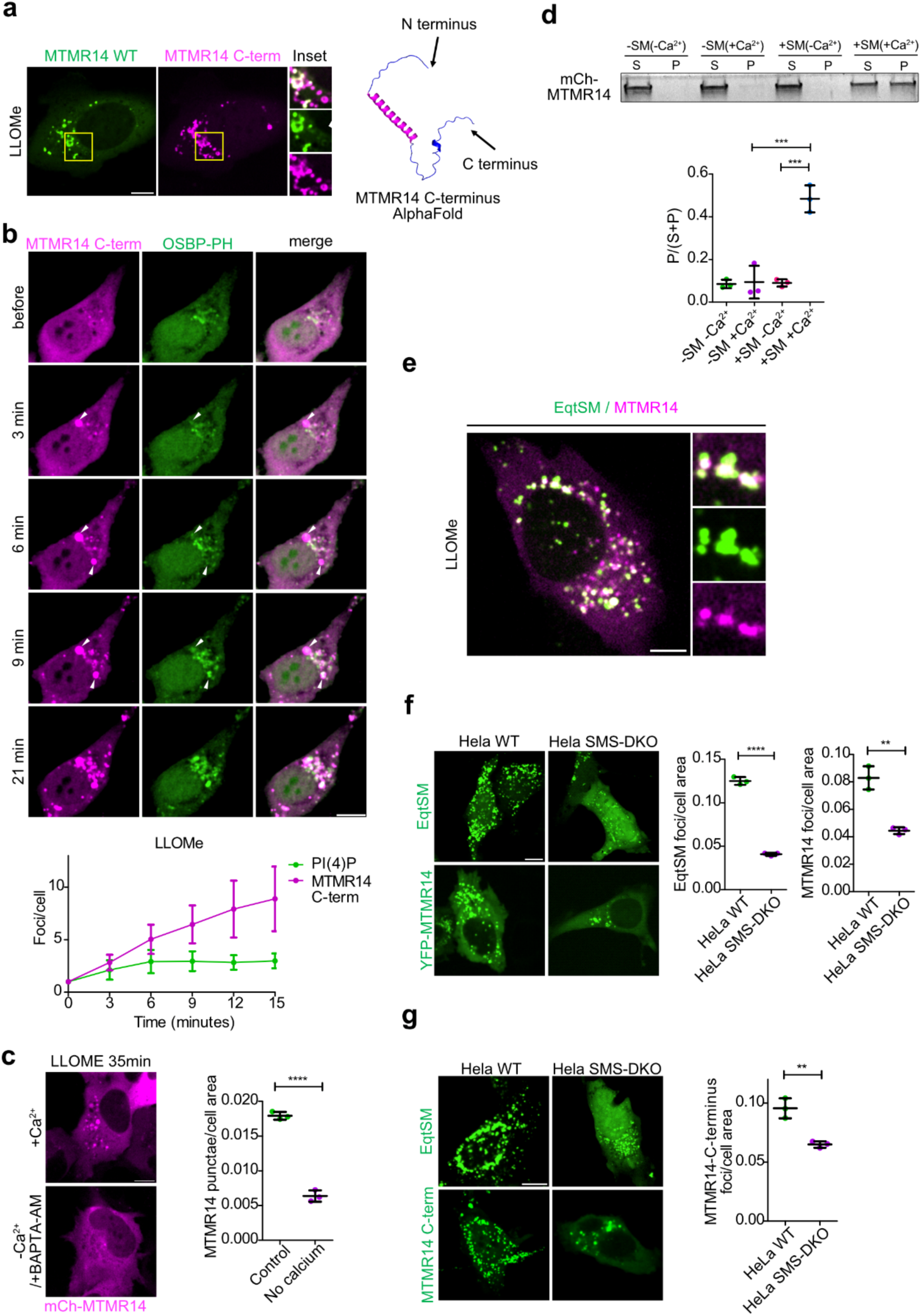
MTMR14 recruitment to damaged lysosomes requires cytosolic calcium and sphingomyelin. (a) The isolated MTMR14 C-terminal tail is recruited to damaged lysosomes. Left: U2OS cells co-expressing full length YFP-MTMR14 and mCherry-MTMR14 C-terminus imaged after 2 h 1mM LLOMe treatment. Scale bar, 10 µm. Right: schematic illustration of the MTMR14 C-terminus predicted by Alpha Fold 2. (b) Top: representative confocal live cell images of 0.5mM LLOMe-induced recruitment kinetics of mCherry-MTMR14 C-terminus and PI(4)P sensor GFP-OSBP-PH co-expressed in HeLa cells. MTMR14 C-terminus foci preceded by GFP-OSBP PH recruitment are indicated by white arrowheads. Bottom: LLOMe-induced recruitment kinetics of mCherry-MTMR14 C-terminus and PI(4)P sensor GFP-OSBP-PH co-expressed in HeLa cells quantified from time lapse live cell imaging and plotted as fold change over untreated. n = 3 independent experiments from 32 cells analyzed in total. Data are mean ± s.e.m. (c) Ca^2+^ dependence of MTMR14 recruitment to lysosomes. Left: representative live cell confocal images of U2OS cells inducibly expressing mCherry-MTMR14 35 minutes post-1mM LLOMe treatment in Ca^2+^ containing media (DMEM/FBS) or Ca^2+^ depleted media (2 mM EDTA in DMEM/FBS + 100 µM BAPTA-AM). Scale bar, 10 µm. Right: quantification of MTMR14 puncta/cell area in images as shown on the left. t test (n = 3 independent experiments, total number of cells is 286 for control and 192 for calcium depleted conditions). (d) Top: representative coomassie blue staining of supernatant and pellet fractions of purified mCherry-MTMR14 incubated with control and sphingomyelin-loaded liposomes in presence and absence of calcium. Bottom: densitometric analysis of binding of purified mCherry-MTMR14 to liposomes from gel images as shown in the top panel. one-way ANOVA (n = 3 independent experiments). SM, sphingomyelin; S, supernatant; P, pellet. (e) Representative live cell confocal image of 1 h 1mM LLOMe-treated U2OS cells co-expressing mCherry-MTMR14 and eGFP-EqtSM. Scale bar, 10 µm. (f) Left: representative live cell confocal images of 1 h 1mM LLOMe-treated WT or SMS-DKO HeLa cells expressing eGFP-EqtSM or YFP-MTMR14. Scale bar, 10 µm. Middle: quantification of the number of eGFP-EqtSM puncta/cell area in WT and SMS-DKO HeLa cells. t test (n = 3 independent experiments, total number of cells is 171 for WT and 172 for SMS-DKO). Right: quantification of the number of MTMR14 puncta/cell area in WT and SMS-DKO HeLa cells. t test (n = 3 independent experiments, total number of cells is 104 for WT and 117 for SMS-DKO). (g) Left: representative live cell confocal images of 1 h 1mM LLOMe-treated WT or SMS-DKO HeLa cells expressing eGFP-EqtSM or mCherry-MTMR14 C-terminal alpha helix. Scale bar, 10 µm. Right: quantification of the number of MTMR14 C-terminal alpha helix puncta/cell area in WT and SMS1/2 DKO 1 h 1mM LLOMe-treated HeLa cells. t test (n = 3 independent experiments, total number of cells is 166 for WT and 107 for SMS-DKO). Statistical analyses were performed using GraphPad Prism. Two-tailed unpaired t-test, paired t-test or one-sample t-tests were conducted using column statistics to compare the sample means to a hypothetical value of 1 or one-way ANOVA with Tukey’s multiple comparisons test. All bar graphs represent mean ± SD unless otherwise stated. ***p < 0.001, **p < 0.01, *p < 0.05. See also Extended Data Fig. 3.

Second, we aimed to determine the trigger that recruits MTMR14 to damaged lysosomes. A prominent feature of lysosomal membrane damage induced by diverse agents is the increased leakage of calcium ions into the cytoplasm of cells (Extended Data Figure 3b). Consistently, we found that growth in calcium-free medium and chelation of cytosolic calcium by BAPTA-AM (Extended Data Figure 3c) occluded MTMR14 recruitment to lysosomes damaged by LLOMe (Figure 2c). Further analysis revealed MTMR14 to be recruited to membranes in response to damage induced by LLOMe, GPN (i.e. lysosomes), BAC, and MSDH (i.e. to plasma membrane puncta) but not upon increase of cytosolic calcium in the presence of ionomycin (Extended Data Figure 3d). This suggests that calcium is required but not sufficient for MTMR14 recruitment to lysosomes. To identify the missing factor involved in MTMR14 recruitment to damaged membranes, we purified recombinant mCherry-tagged MTMR14 from overexpressing Expi293F cells (Extended Data Fig. 3e) and assayed its ability to associate with liposomes. We found MTMR14 to potently bind to liposomal membranes containing Sphingomyelin in the presence of calcium (Figure 2d). Sphingomyelin is a lipid enriched on the lumenal leaflet of the lysosomal limiting membrane and previous data have shown that lysosomal membrane damage triggers calcium-activated Sphingomyelin scrambling at lysosomes^23^. Consistently, we found MTMR14 to colocalize with Sphingomyelin monitored by eGFP-tagged Equinatoxin II at damaged lysosomes (Figure 2e). Furthermore, depletion of sphingomyelin and the concomitant accumulation of hexosylceramides in sphingomyelin synthase 1/2 double KO (DKO) U2OS (Extended Data Fig. 3f,g) markedly reduced recruitment of full-length MTMR14 or its C-terminal domain to damaged lysosomes *in vivo* (Figure 2f,g) with a similar effect size as Equinatoxin II (Extended Data Fig. 3h).

These findings demonstrate that MTMR14 is rapidly (i.e. within ≤ 3 min) recruited to damaged lysosomes by direct calcium-triggered association with Sphingomyelin to erase PI(3)P.

## MTMR14 triggers a local signaling lipid switch to aid lysosomal membrane repair

To mechanistically dissect the functional role of MTMR14 in the cellular response to lysosomal membrane damage, we used a well-established method that probes lysosomal membrane integrity based on lysotracker, i.e. a vital dye which selectively stains intact acidic endolysosomal compartments. This assay monitors the ability of cells to repair damaged lysosomes upon acute damage by pulsed treatment with LLOMe followed by washout to enable restoration of the membrane permeability barrier. MTMR14 KO cells suffered from a defect in the recovery of lysosomal membrane integrity (Figure 3a). A second independent MTMR14 KO line (Extended Data Fig. 2e) displayed a very similar phenotype (Extended Data Fig. 4a). Recovery of lysotracker was unaffected by KO of MTM1 (Extended Data Fig. 4b), a closely related endolysomal PI(3)P phosphatase only mildly recruited to damaged lysosomes (Extended Data Fig. 1e). MTMR14 KO cells also suffered from impaired recovery of lysosomal membrane integrity following chronic sustained exposure to LLOMe (Figure 3b, Extended Data Fig. 4c), i.e. a protocol that may mimic to some degree pathophysiological conditions induced by the sustained presence of non-degradable agents such as tau protein aggregates. Consistently, defective recovery of lysosome function was also overt from the persistent accumulation of Galectin3 in MTMR14 KO compared to wild-type U2OS cells overexpressing neuropathogenic tau (P301L) (Figure 3c). Defective lysosomal membrane repair was paralleled by elevated death of MTMR14 KO U2OS cells (Figure 3d) or MTMR14-depleted C2C12 myoblasts (Extended Data Fig. 4d,e) subjected to lysosomal membrane damage.

**Figure 3.**
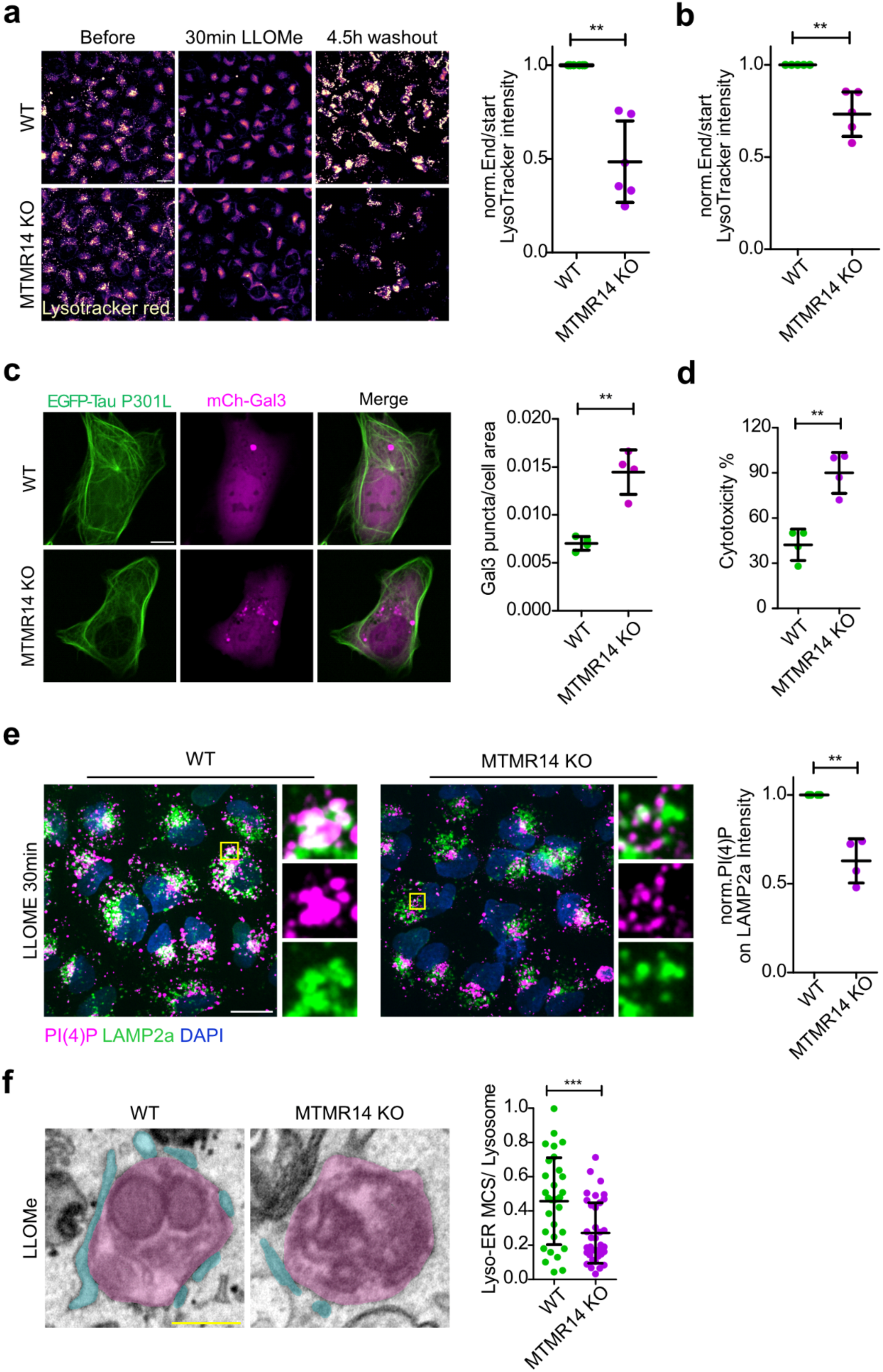
MTMR14 is required for cellular adaptation in response to LMP. (a) Left: defective lysosomal recovery in MTMR14 KO cells. Representative confocal images of LysoTracker loaded WT and MTMR14 KO clone #1 U2OS cells imaged before LLOMe treatment, after 30 minutes LLOMe treatment, and 4.5 h after washout of LLOMe. Scale bar, 20 µm. Right: quantification of mean LysoTracker intensity/field of view, fold change over control. t test (n = 6 independent experiments, each datapoint represents 10 fields of view, one field of view containing 35-50 cells, with a size of 3328 × 3328 µm). (b) Quantification of mean LysoTracker intensity/field of view after 5 h chronic LLOMe treatment in U2OS MTMR14 KO clone #1 fold change over WT U2OS cells, t test (n = 5 independent experiments, each datapoint represents 10 fields of view, one field of view containing 35-50 cells, with a size of 3328 × 3328 µm). (c) Left: Loss of MTMR14 causes elevated lysosomal membrane damage monitored by mCherry-Galectin3 in U2OS cells co-expressing mutant eGFP-Tau (P301L) for 24 hours. Confocal images of representative cells. Scale bar, 10 µm. Right: Quantification of mCherry-Galectin3 puncta/cell area in U2OS WT and MTMR14 KO cells after co-expressing mutant eGFP-Tau (P301L) for 24 hours. t test (n = 4 independent experiments, total number of cells is 107 for both WT and MTMR14 KO). (d) Quantification of cytotoxicity using LDH assay in 5 h 4mM LLOMe-treated WT and MTMR14 KO U2OS cells. t test (n = 4 independent experiments, each datapoint represents triplicate measurements). (e) MTMR14 regulates PI(4)P generation in response to LMP. Left: representative confocal images of WT and MTMR14 KO U2OS cells stained with DAPI and antibodies against PI(4)P and Lamp2a in control conditions (DMSO) and after 30 minutes LLOMe treatment. Right: quantification of PI(4)P intensity on Lamp2a detections in images as shown in the left panel. t test (n = 4 independent experiments, total number of fields of view is 109 for WT and 114 for MTMR14 KO, one field of view containing 10-15 cells, with a size of 1664 × 1664 µm). (f) Left: representative colored electron micrographs of lysosomes in WT and MTMR14 KO U2OS cells treated with LLOMe, ER is colored in blue and lysosomes in pink. Scale bar, 200 nm. Right: length of lysosome-ER MCS relative to lysosomal perimeter in WT and MTMR14 KO U2OS cells. t test, WT (n = 31 lysosomes), MTMR14 KO (n = 37 lysosomes). Statistical analyses were performed using GraphPad Prism. Two-tailed unpaired t-test, paired t-test or one-sample t-tests were conducted using column statistics to compare the sample means to a hypothetical value of 1 or one-way ANOVA with Tukey’s multiple comparisons test. All bar graphs represent mean ± SD unless otherwise stated. ***p < 0.001, **p < 0.01, *p < 0.05. See also Extended Data Fig. 4.

Our findings suggest that MTMR14 may be important for the recovery of functional lysosomes, e.g. by preserving or restoring the integrity of lysosomal membranes. To test this in a pathophysiological setting, we exposed WT and MTMR14 KO cells to a *ΔsifA* mutant strain of the enteropathogen *Salmonella enterica Typhimurium,* which predominantly replicates in the cytoplasm following escape from the endolysosomal PI(3)P-enriched *Salmonella*-containing vacuole (SCV)^56^. As predicted, replication of the mutant strain was increased in the absence of MTMR14 (Extended Data Fig. 4f,g), suggesting that MTMR14 loss renders the replicative vacuole (i.e. a specialized lysosome) more fragile to facilitate bacterial escape and replication in the cytoplasm.

We hypothesized that the role of MTMR14 in preserving and/ or restoring the integrity of lysosomal membranes following damage or bacterial infection, may relate to a switch in lysosomal signaling lipid identity that involves PI(3)P hydrolysis on lysosomes via MTMR14 to facilitate lysosomal PI(4)P synthesis and, thereby, membrane repair, akin to the lysosomal starvation response ^47^. To test this, we monitored the effects of MTMR14 loss on lysosomal PI(4)P synthesis in response to LMP. We observed that KO of MTMR14 from U2OS cells markedly reduced the damage-induced synthesis of PI(4)P on lysosomes observed in control cells (Figure 3e, Extended Data Fig. 4h). As lysosomal PI(4)P is of key importance for lysosomal repair by ER-based MCS via the PITT pathway^22^, we conducted quantitative morphometric analysis of ER/ lysosome MCS by electron microscopy. We found a significant reduction of ER/ lysosome MCS in MTMR14 KO compared to wild-type cells treated with LLOMe (Figure 3f). In contrast, loss of MTMR14 in KO cells or the near complete depletion of PI(3)P on lysosomes upon inhibition of VPS34 PI 3-kinase in the presence of VPS34-IN1 (consistent with ^18, 19^) did not affect the recruitment of ESCRT proteins to damaged lysosomes (Extended Data Fig. 5a-e), consistent with ^18^.

Collectively, these results unravel an MTMR14-based lysosomal signaling lipid switch that aids in preserving and/ or restoring the integrity of lysosomal membranes following damage.

## PI(3)P depletion by MTMR14 represses canonical mTORC1 signaling

Previous data suggest that lysosomal membrane damage inhibits mTORC1 signaling at lysosomes ^34^ and we and others identified PI(3)P as a potent regulator of canonical mTORC1 signaling, including the downstream regulation of PI(4)P kinase PI4K2A^47, 57^ (Figure 4a). We therefore hypothesized that the damage-induced depletion of lysosomal PI(3)P via MTMR14 (see Figure 1) may coordinate PI(4)P-mediated repair with metabolism by controling mTORC1 signaling. Interestingly, we found *MTMR14* expression to be inversely correlated with body weight and body mass index (Extended Data Fig. 6a,b), while MTMR14 KO mice suffer from obesity and metabolic dysregulation ^45^, suggesting a role in metabolic control. To directly probe the potential function of MTMR14 in the control of mTORC1 signaling in cellular response to LMP, we analyzed the phosphorylation status of the canonical downstream substrate p70 S6 kinase (S6K) in U2OS cells subjected to lysosomal membrane damage. S6K phosphorylation was prominently repressed upon LMP (Figure 4b), consistent with ^34^. Strikingly, damage-induced repression of mTORC1-S6K signaling was occluded by KO of MTMR14 (Figure 4b). mTORC1 downstream signaling pathways could affect adaptation to LMP via multiple mechanisms such as lysophagy, the PITT pathway, translation regulation, or lysosome biogenesis (Figure 4a). We have recently shown that in nutrient replete conditions S6K inhibits the central PITT regulator and PI(4)P synthesizing enzyme PI4K2A^32^, suggesting that LMP could facilitate PI(4)P generation via inhibition of mTORC1-S6K signaling (Figure 4a). Indeed, LLOMe led to decreased PI4K2A phosphorylation (Extended Data Fig. 6c) and loss of MTMR14 occluded repression of S6K activity towards its canonical substrate S6 protein (Figure 4c; Extended Data Fig. 6d,e). Similarly, MTMR14 loss impacted phosphorylation of ULK1 kinase (Extended Data Fig. 6f), a major regulator of autophagy, and of the translation regulatory factor 4E-BP1 (Extended Data Fig. 6g). In contrast, non-canonical mTORC1 signaling via TFEB was unaffected by KO of MTMR14 or acute repression of PI(3)P synthesis in the presence of VPS34-IN1 (Extended Data Fig. 6h-j). Importantly, defective repression of LLOMe-induced inhibition of canonical mTORC1-S6K-S6 signaling in MTMR14 KO cells was rescued by re-expression of active MTMR14 (Figure 4d,e; Extended Data Fig. 6g,k) but not by a phosphatase-inactive mutant of MTMR14 (Figure 4e, Extended Data Fig. 6k). Hence, active MTMR14, likely by hydrolyzing PI(3)P, is required for the LMP-induced repression of canonical mTORC1-S6K signaling.

**Figure 4.**
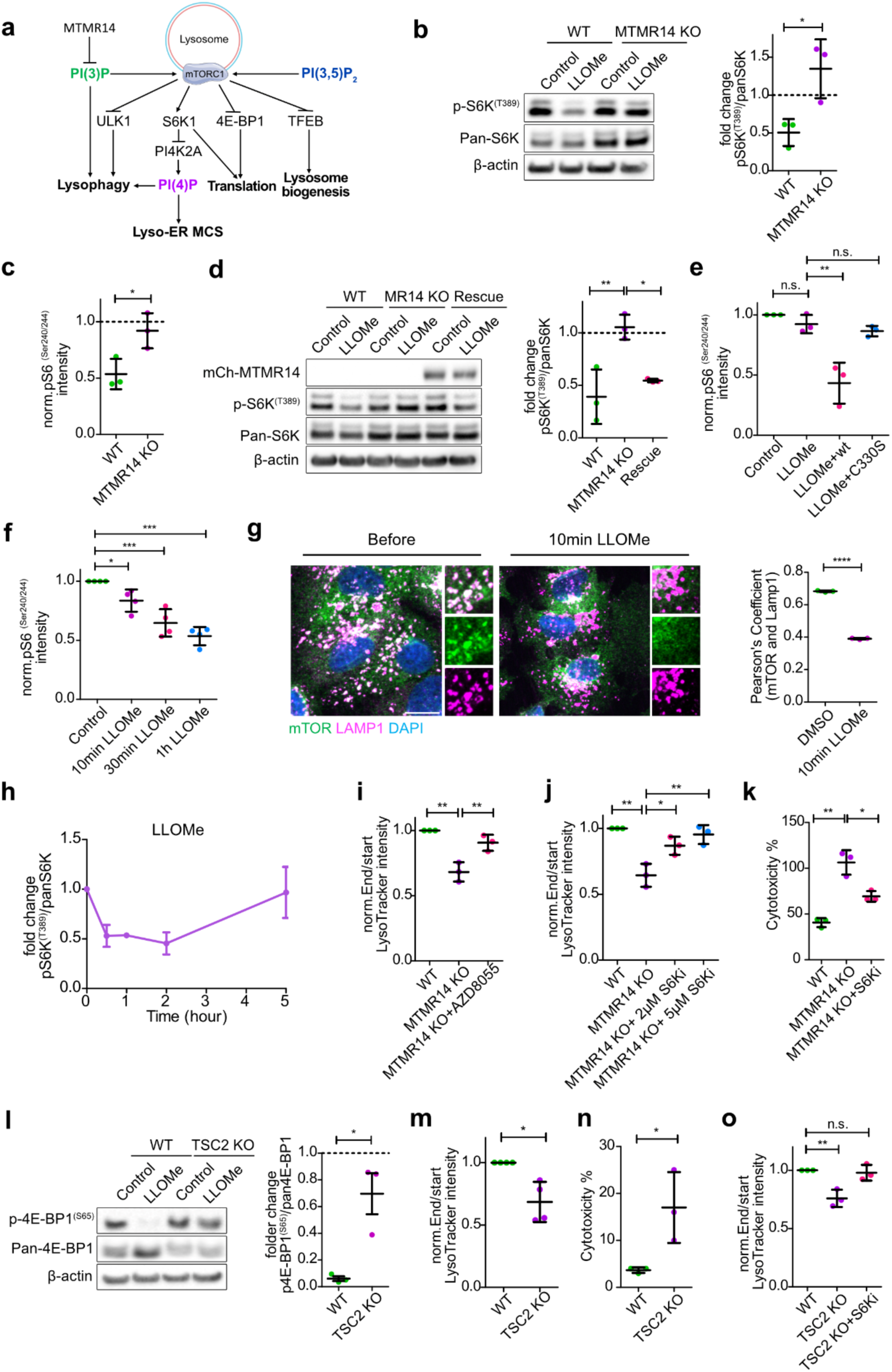
MTMR14-dependent inhibition of mTORC1 is required for recovery from LMP. (a) Schematic illustration of phosphoinositide pathways impinging on mTORC1 and their possible effects on LMP adaptation processes. (b) Inhibition of mTORC1 in response to LLOMe is dependent on MTMR14. Left: Representative immunoblot analysis of pS6K T389/ total S6K levels in control or 2 h LLOMe-treated WT and MTMR14 KO U2OS cells. Right: quantification of representative data from n = 3 independent experiments (t test). Dotted line denotes pS6K/ total S6K levels in DMSO controls set to 1. (c) Quantification of pS6 S240/244 intensity by immunofluorescence in WT and MTMR14 KO U2OS cells upon 1h LLOMe treatment. n = 3 independent experiments (t test), each datapoint represents 15 fields of view. Dotted line denotes pS6 intensity in DMSO controls set to 1. Representative confocal images are shown in Extended Data Fig. 6e. (d) MTMR14 re-expression in KO cells rescues decreased S6K phosphorylation upon LLOMe treatment. Left: Representative immunoblot analysis of U2OS WT, MTMR14 KO, and MTMR14 KO cells expressing mCherry-MTMR14 under doxycycline control subjected to 2 h DMSO control or LLOMe treatment. Right: quantification of immunoblots from n = 3 independent experiments. One-way ANOVA. Dotted line denotes pS6K T389/ total S6K levels in DMSO controls set to 1. (e) Expression of MTMR14 WT but not of catalytically inactive MTMR14 C330S in MTMR14 KO U2OS cells rescues decreased S6 phosphorylation upon LLOMe treatment. Quantification of pS6 S240/244 intensity by immunofluorescence in MTMR14 KO U2OS cells in different conditions. one-way ANOVA (n = 3 independent experiments), control 105 cells, LLOMe 116 cells, LLOMe + MTMR14 WT 75 cells, LLOMe + MTMR14 C330S 96 cells. Representative confocal images are shown in Extended Data Fig. 6k. (f) Quantification of pS6 S240/244 intensity by immunofluorescence in U2OS cells upon treatment with LLOMe for the indicated times. one-way ANOVA, each datapoint represents 15 fields of view. Representative confocal images are shown in Extended Data Fig. 6l. (g) Left: representative confocal images of 10 minutes control (DMSO) and LLOMe-treated U2OS cells stained with DAPI and antibodies against mTOR and LAMP1. Scale bar, 10 µm. Right: quantification of Pearson’s Coefficient between mTOR and LAMP1. t test (n = 3 independent experiments, each datapoint represents 15 fields of view, one field of view containing 10-20 cells, with a size of 1664 × 1664 µm). (h) Quantification of pS6K T389/ total S6K levels in U2OS cells treated with LLOMe for the indicated times and analyzed by immunoblotting, n = 3 independent experiments. Data are mean ± s.e.m. Representative immunoblot is shown in Extended Data Fig. 6m. (i) Quantification of mean LysoTracker intensity/field of view fold change over control after 5 h LLOMe treatment in U2OS WT, MTMR14 KO, or MTMR14 KO cells treated with 200 nM AZD8055. One-way ANOVA (n = 3 independent experiments, each datapoint represents 10 fields of view, one field of view containing 35-50 cells, with a size of 3328 × 3328 µm). (j) S6K inhibition partially rescues lysosomal recovery defects in LLOMe-exposed MTMR14 KO cells. Lysotracker recovery measured as fold change over control in 5 h LLOMe-treated WT vs MTMR14 KO U2OS cells in the presence of the indicated concentrations of S6K inhibitor (S6Ki) from n = 3 independent experiments, one-way ANOVA, each datapoint represents 10 fields of view, one field of view containing 35-50 cells, with a size of 3328 × 3328 µm. (k) S6K inhibition partially rescues cytotoxicity in LLOMe-exposed MTMR14 KO cells. Cytotoxicity was measured by LDH assay in U2OS WT, MTMR14 KO, and MTMR14 KO cells treated with 2 µM S6Ki. One-way ANOVA (n = 3 independent experiments, each datapoint represents triplicate measurements). (l) TSC2 KO cells fail to inhibit mTORC1 in response to LMP. Left: representative immunoblot of p4E-BP1 S65 phosphorylation in HeLa WT and TSC2 KO cells treated with DMSO control or LLOMe. Right: quantification of immunoblots as shown in the left panel. t test (n = 3 independent experiments). Dotted line denotes p4E-BP1 S65/ total 4E-BP in controls set to 1. (m) TSC2 KO cells show defects in lysosome recovery upon exposure to LLOMe. Quantification of mean LysoTracker intensity/field of view fold change over control after 5 h LLOMe treatment in HeLa WT, TSC2 KO, and TSC2 KO cells treated with 5 µM S6Ki. one-way ANOVA (n = 3 independent experiments, each datapoint represents 10 fields of view, one field of view containing 35-50 cells, with a size of 3328 × 3328 µm). (n) TSC2 KO cells display increased cell death upon exposure to LLOMe. Cytotoxicity in HeLa WT and TSC2 KO cells after 5 h 4mM LLOMe treatment, t test (n = 4 independent experiments, each datapoint represents triplicate LDH assay measurements). (o) Lysosome recovery defects in TSC2 KO cells are rescued by S6K inhibition. Lysotracker recovery measured as fold change over control in 5 h LLOMe-treated WT vs TSC2 KO HeLa cells in the presence of control (DMSO) or S6K inhibitor (S6Ki) from n = 3 independent experiments, one-way ANOVA, each datapoint represents 10 fields of view, one field of view containing 35-50 cells, with a size of 3328 × 3328 µm. Statistical analyses were performed using GraphPad Prism. Two-tailed unpaired t-test, paired t-test or one-sample t-tests were conducted using column statistics to compare the sample means to a hypothetical value of 1 or one-way ANOVA with Tukey’s multiple comparisons test. All bar graphs represent mean ± SD unless otherwise stated. ***p < 0.001, **p < 0.01, *p < 0.05. See also Extended Data Fig. 6.

To unravel a potential causal relationship between LMP-induced PI(3)P depletion by MTMR14 (see Figures 1 and 2) and the repression of canonical mTORC1 signaling, we monitored the kinetics of S6K and ribosomal S6 protein phosphorylation during the damage response. Confocal imaging revealed that reduced ribosomal S6 protein phosphorylation was detectable as early as 10 min post-LMP (i.e. about 5 min after the onset of MTMR14 recruitment, see Figure 1), reached its low steady-state levels by about 1h (Figure 4f, Extended Data Fig. 6l), and was accompanied by the partial dissociation of mTOR kinase from lysosomes marked by LAMP1 (Figure 4g). The initial repression of canonical mTORC1-S6K signaling post-LMP was followed by slow recovery of nutrient signaling over many hours as damaged lysosomal membranes were repaired or removed (Figure 4h) in various cell types (Extended Data Fig. 6m-o).

Collectively, these data suggest that repression of canonical mTORC1-S6K signaling via MTMR14-mediated hydrolysis of PI(3)P facilitates the recovery of lysosome function during membrane damage (Figure 4a). To test this hypothesis, we first analyzed the effects of repressing mTORC1-S6K signaling in cells lacking MTMR14. Indeed, we found that pharmacological inhibition of mTOR-S6K signaling (Extended Data Fig. 6p) facilitated recovery of lysosome function monitored by lysotracker (Figure 4i,j) and ameliorated LLOMe-induced cell death in MTMR14 KO cells (Figure 4k). Second, we monitored the susceptibility of cells lacking TSC2, a well-established repressor of canonical mTORC1-S6K signaling to LMP. Loss of TSC2 occluded the LLOMe-induced repression of canonical mTORC1 signaling, akin to KO of MTMR14 (Figure 4l), resulting in impaired recovery of lysosomal membrane integrity and induction of cell death (Figure 4m,n). Pharmacological inhibition of S6K signaling rescued defective lysosomal membrane repair in TSC2 KO cells (Figure 4o).

These data indicate that repression of canonical mTORC1-S6K signaling via MTMR14-mediated hydrolysis of PI(3)P facilitates the recovery of lysosome function during membrane damage.

## MTMR14-mediated repression of canonical mTORC1 signaling blocks protein translation to aid cell survival

How does repression of canonical mTORC1 signaling via MTMR14-mediated PI(3)P hydrolysis facilitate the recovery of lysosome function and normal cell physiology? Previous studies and our work described thus far have demonstrated that repression of mTORC1-S6K signaling (i.e. via PI(3)P hydrolysis on lysosomes) inhibits lysosomal PI(4)P synthesis and ER-lysosome MCS formation by inactivation of PI4K2A^47^ (compare Figure 4a), a pathway that can be induced by LMP (Figure 3). Second, work by us^47^ and others^31^ has shown that lysosomal PI(3)P promotes mTORC1 activation and, thereby, may facilitate protein translation, i.e. via S6K activation of ribosomal S6 protein^58^ and hyperphosphorylation of 4E-BPs to promote the recruitment of mRNAs to ribosomes. Based on these data and the observation that loss of lysosomal membrane integrity impairs proteostasis^59, 60^, we hypothesized that damage-induced PI(3)P depletion via MTMR14 may repress protein translation. We probed this hypothesis by analyzing the impact of LMP on protein synthesis. LLOMe-induced damage potently repressed protein translation monitored by puromycin incorporation into nascent chains (Figure 5a, Extended Data Fig. 7a,b). In contrast, MTMR14 KO cells failed to repress protein synthesis in response to lysosomal membrane damage (Figure 5b,c). Importantly, pharmacological inhibition of S6K restored defective LLOMe-induced repression of protein translation in MTMR14 KO cells (Figure 5d). These data suggest that mTORC1-S6K-dependent protein synthesis is controlled by PI(3)P via MTMR14.

**Figure 5.**
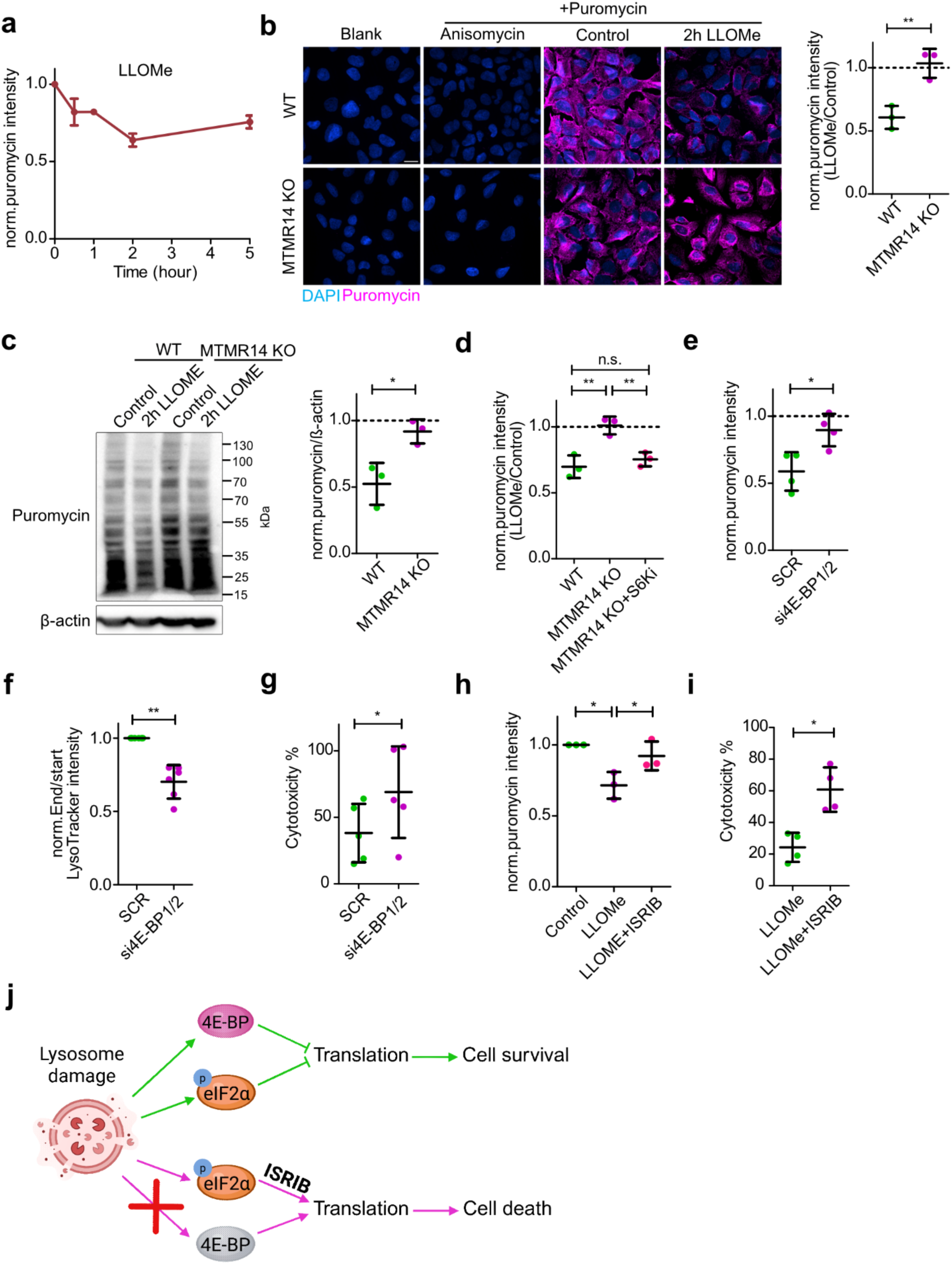
Repression of translation safeguards cell survival in response to LMP. (a) Time course of protein translation repression in LLOMe-treated U2OS cells analyzed by puromycin staining, n = 4 independent experiments, each datapoint represents 20 fields of view, one field of view containing 10-20 cells, with a size of 1664 × 1664 µm. Data are mean ± s.e.m. Representative confocal images are shown in Extended Data Fig. 7a. (b) Left: representative confocal images of WT and MTMR14 KO U2OS cells treated as indicated and stained for puromycin and DAPI. Scale bar, 20 µm. Right: quantification of images as shown in the left panel, t test (n = 3 independent experiments, each datapoint represents 20 fields of view, one field of view containing 10-20 cells, with a size of 1664 × 1664 µm). Dotted line denotes puromycin intensity in DMSO controls set to 1. (c) Translation repression in response to LMP is impaired in MTMR14 KO cells. Left: representative immunoblot of control and LLOMe-treated WT and MTMR14 KO U2OS cell lysates. Right: densitometric quantification of blots as shown in the left panel. t test (n = 3 independent experiments). Dotted line denotes puromycin intensity in DMSO controls set to 1. (d) Analysis as in (b) of WT, MTMR14 KO, and MTMR14 KO U2OS cells treated with S6K inhibitor (S6Ki) stained for puromycin and DAPI. one-way ANOVA (n = 3 independent experiments, each datapoint represents 20 fields of view, one field of view containing 10-20 cells, with a size of 1664 × 1664 µm). Dotted line denotes puromycin intensity in DMSO controls set to 1. (e) Knockdown of 4E-BP1 increases translation. Puromycin intensity quantification in U2OS cells transfected with scr control or siRNA against 4E-BP1 and 4E-BP2 (si4E-BP1/2), treated with LLOMe for 2 h, and stained with Puromycin and analysed by confocal imaging. t test (n = 4 independent experiments, each datapoint represents 15 fields of view, one field of view containing 10-20 cells, with a size of 1664 × 1664 µm). Dotted line denotes puromycin intensity in DMSO controls set to 1. (f) 4E-BP knockdown prevents full lysosome recovery after LLOMe treatment. LysoTracker recovery measured as fold change over control in 5 h LLOMe-treated U2OS cells transfected with scr control or siRNA against 4E-BP1 and 4E-BP2 (si4E-BP1/2). t test (n = 5 independent experiments, each datapoint represents 10 fields of view, one field of view containing 35-50 cells, with a size of 3328 × 3328 µm). (g) Knockdown of translation repressor 4E-BP increases susceptibility to cell death in response to LLOMe. U2OS cells transfected with scr control vs siRNA against 4E-BP1 and 4E-BP2 (si4E-BP1/2) were treated with 4mM LLOMe for 5 h and cytotoxicity was measured with LDH assay, t test (n = 5 independent experiments performed in triplicate or duplicate measurements). (h) Analysis as in (b) of control (DMSO), LLOMe, and LLOMe plus 500 nM ISRIB treatment in U2OS cells stained for puromycin and DAPI. One-way ANOVA (n = 3 independent experiments, each datapoint represents 15 fields of view, one field of view containing 10-20 cells, with a size of 1664 × 1664 µm). (i) Translation disinhibition by ISRIB increases susceptibility to cell death in response to LLOMe. Cytotoxicity measurement by LDH assay in U2OS cells treated with control (DMSO), 4mM LLOMe, or LLOMe plus 500 nM ISRIB for 5 h, t test (n = 4 independent experiments). (j) Schematic illustration showing that ISRIB treatment or 4E-BP silencing prevents downregulation of translation, leading to cell death. Statistical analyses were performed using GraphPad Prism. Two-tailed unpaired t-test, paired t-test or one-sample t-tests were conducted using column statistics to compare the sample means to a hypothetical value of 1 or one-way ANOVA with Tukey’s multiple comparisons test. All bar graphs represent mean ± SD unless otherwise stated. ***p < 0.001, **p < 0.01, *p < 0.05. See also Extended Data Fig. 7.

A common pattern shared across stress responses is decreased global protein synthesis, thereby lowering the overall proteostatic burden, while still synthesizing select proteins needed to resolve the stress^37-39^. We therefore reasoned that MTMR14-mediated repression of mTORC1-dependent protein synthesis may aid the recovery of lysosome function and cell survival by lowering the proteostatic burden. We tested this notion via independent genetic and pharmacological manipulations. First, we depleted cells of the mTORC1-controlled translational repressor 4E-BP1/2. Cells depleted of 4E-BP1/2 failed to repress protein translation in response to LMP (Figure 5e) and were defective with respect to their ability to recover lysosomal membrane integrity (Figure 5f), resulting in increased cell death (Figure 5g), e.g. phenotypes similar to KO of MTMR14 (compare Figures 3; 5b,c). Second, we applied Integrated Stress Response Inhibitor (ISRIB), a small molecule that reverses the repressive effects of the integrated stress response on protein synthesis by overcoming the effects of eIF2a phosphorylation^61^. Application of ISRIB to LLOMe-treated cells significantly rescued reduced global protein synthesis (Figure 5h) and, importantly, aggravated LMP-triggered cell death (Figure 5i).

Collectively, these results suggest a model according to which repression of protein synthesis via MTMR14-mediated inhibition of canonical mTORC1 signaling facilitates recovery of lysosome function and survival of cells subjected to lysosomal membrane damage (Figure 5j).

## MTMR14-mediated repression of canonical mTORC1 signaling enables induction of lysophagy

Lysosomal membrane damage, in addition to repair processes, induces lysophagy. While induction of lysophagy is known to require repression of canonical mTORC1 signaling via ULK1^30^, possibly via PI(3)P hydrolysis, the clearance of damaged lysosomes by autophagy depends on PI(4)P synthesis via PI4K2A^43, 47, 62^ as well as on PI(3)P made by VPS34^63^ (as further shown below). To kinetically resolve this process and its potential regulation by PI(3)P turnover and synthesis, we monitored the time course of lysophagy induction via the lysophagy sensor TMEM192-mKeima^35^ in cells subjected to lysosomal membrane damage. We found lysophagy to be induced with a time delay of about 2 h post-damage in HeLa (Figure 6a, Extended Data Fig. 8a) or U2OS cells (Extended Data Fig. 8b), i.e. at a time point when PI(3)P levels had substantially recovered (see Figure 1b,c). This suggests a model (Figure 6b), in which acute recruitment of MTMR14 within minutes (see Figure 1g) inhibits canonical mTORC1 signaling (Figure 4) to release ULK1 from mTORC1 repression (compare Extended Data Fig. 6f). This enables the induction of lysophagy once PI(3)P levels resurge after about 2 h post-damage induction. We tested this model at several different levels. First, we probed the contribution of lysophagy to the recovery of lysosome function after damage. Pharmacological blockade of lysophagy in the presence of the ULK1 inhibitor MRT68921 impaired the recovery of lysosomal membrane integrity monitored by lysotracker fluorescence in U2OS (Figure 6c) and microglial BV2 cells (Extended Data Fig. 8d), and increased LLOMe-induced death of U2OS cells (Figure 6d), confirming the importance of lysophagy for the recovery of lysosome function. Second, we probed whether MTMR14-mediated repression of canonical mTORC1 signaling is required for lysophagy induction upon damage. Indeed, we found that KO of MTMR14 impaired lysophagy measured by TMEM192-mKeima in U2OS cells (Figure 6e, Extended Data Fig. 8c). This phenotypic defect was largely rescued by mTOR kinase inhibition (Figure 6f), consistent with the hypothesis that MTMR14-mediated PI(3)P hydrolysis serves to release ULK1 from mTORC1 repression. Third, we probed whether LMP-induced lysophagy depends on the resurgence of PI(3)P levels. Blocking PI(3)P synthesis by inhibition of phosphoinositide 3-kinase VPS34 led to impaired lysophagy to a similar degree as ULK1 inhibition (Figure 6g,h) underscoring the important role of PI(3)P in lysophagy. VPS34 inhibition also led to reduced lysotracker recovery after prolonged LLOMe treatment in U2OS or microglial BV2 cells (Figure 6i; Extended Data Fig. 8d). and increased cell death in U2OS (Figure 6j) and BV2 cells (Extended Data Fig. 8e) whereas inhibition of the P(3,5)P_2_ kinase PIKfyve did not affect lysotracker recovery after LLOMe (Extended Data Fig. 8f).

**Figure 6.**
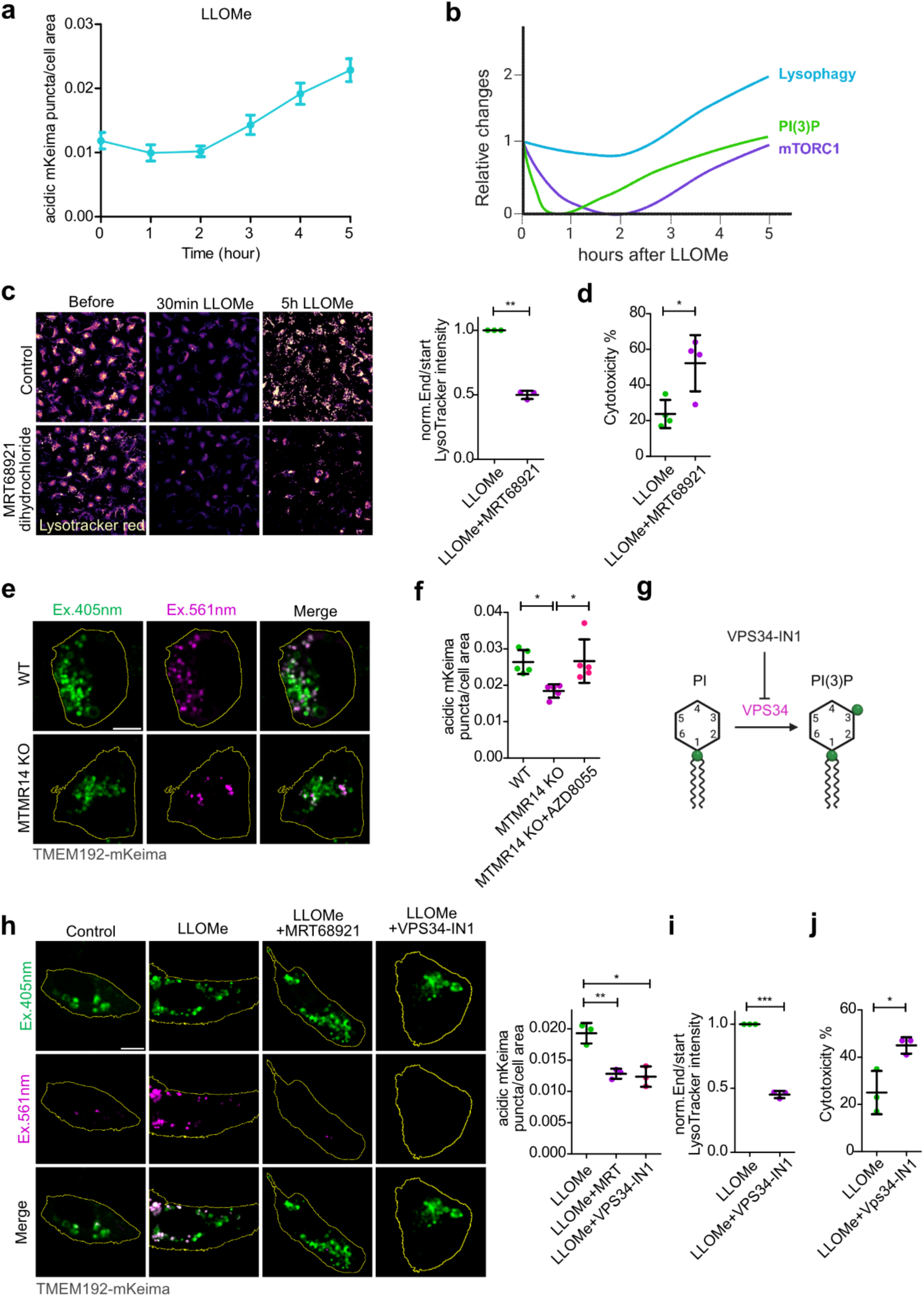
MTMR14-mediated repression of canonical nutrient signalling enables induction of PI(3)P-dependent lysophagy. (a) Time course of LLOMe-induced lysophagy. Quantification of acidic TMEM192-mKeima puncta/cell area in HeLa cells after LLOMe treatment, each datapoint represents 24-50 cells. Data are mean ± s.e.m. Representative confocal images are shown in Extended Data Fig. 8. (b) Schematic illustrating the time course of lysosomal membrane damage-induced changes in lysophagy, PI(3)P, and canonical mTORC1 signalling. Curves were modelled according to experimental data shown in Figure 1c [PI(3)P], Figure 4h (mTORC1), and Figure 6a (Lysophagy). PI(3)P and mTORC1 curves were normalized between 0 and 1 to allow for comparison of the kinetics. (c) Defective lysotracker recovery of U2OS cells treated with ULK1 inhibitor (2µM MRT68921). Left: cells were imaged before and 30min and 5h after LLOMe treatment. Scale bar, 20µm. Right: quantification of representative data shown in left. Mean LysoTracker intensity/field of view, fold change over control. Data are from n = 3 independent experiments (t test), each datapoint represents 10 fields of view, one field of view containing 35-50 cells, with a size of 3328 × 3328 µm. (d) Quantification of cytotoxicity by LDH assay of 5 h 4mM LLOMe-treated U2OS cells in absence or presence of ULK1 inhibitor 2µM MRT68921. t test (n = 4 independent experiments, each datapoint represents 3 replicate measurements). (e) Representative confocal live cell images of LLOMe-treated WT or MTMR14 KO U2OS cells expressing TMEM192-mKeima. Cell outlines are shown in yellow. Quantification is shown in Extended Data Fig. 8c. Scale bar, 10 µm. (f) Quantification of acidic TMEM192-mKeima puncta per cell area in WT or MTMR14 KO U2OS cells treated with LLOMe in the absence or presence of the mTOR inhibitor AZD8055 (200 nM). one-way ANOVA (n = 5 independent experiments, total number of cells is 106 for WT, 106 for, MTMR14 KO and 83 for MTMR14 KO + AZD8055). (g) Schematic illustration showing that VPS34-IN1 inhibits VPS34 leading to PI(3)P depletion. (h) Left: effects of ULK1 and VPS34 inhibition on LLOMe-induced lysophagy in U2OS cells analyzed by confocal live cell imaging of TMEM192-mKeima. Cell outlines are shown in yellow. Scale bar, 10 µm. Right: quantification of acidic mKeima puncta /cell area. One-way ANOVA (n = 3 independent experiments, total number of cells is 76 for LLOMe, 77 for LLOMe+MRT68921, and 78 for LLOMe +VPS34-IN1). (i) Lysotracker recovery measured as fold change over control of 5 h 1mM LLOMe-treated U2OS cells in the absence or presence of 5µM VPS34-IN1. t test (n = 3 independent experiments, each datapoint represents 10 fields of view, one field of view containing 35-50 cells, with a size of 3328 × 3328 µm). (j) Quantification of cytotoxicity of 5 h 4mM LLOMe-treated U2OS cells in absence or presence of 5µM VPS34-IN1 inhibitor. t test (n = 3 independent experiments, each datapoint represents triplicate measurements). Statistical analyses were performed using GraphPad Prism. Two-tailed unpaired t-test, paired t-test or one-sample t-tests were conducted using column statistics to compare the sample means to a hypothetical value of 1 or one-way ANOVA with Tukey’s multiple comparisons test. All bar graphs represent mean ± SD unless otherwise stated. ***p < 0.001, **p < 0.01, *p < 0.05. See also Extended Data Fig. 8.

Together, these data suggest that timed regulation of PI(3)P repression via MTMR14 and resurgence is required for the coordination of lysophagy and lysosomal recovery in response to LMP.

## Ubiquitin-mediated turnover of MTMR14 enables recovery of lysosome function

The findings thus far suggest that lysosomal membrane damage induces a timed series of repair and removal processes that are linked to cellular proteostasis (i.e. downregulation of global protein synthesis) by the transient local repression of canonical mTORC1 signaling, which involves the recruitment of the PI(3)P phosphatase MTMR14 to damaged lysosomes. Our data (Figure 7a) are most compatible with a model, in which LMP first triggers a rapid response within minutes aimed to repair damaged membranes, e.g. via recruitment of ESCRT proteins and lipid transfer via the PITT pathway that is aided by a MTMR14-mediated lysosomal signaling lipid switch via hydrolysis of PI(3)P at damaged lysosomes (compare Figures 1-3). The rapid initial decline of lysosomal PI(3)P mediated by MTMR14 with some delay (10-60 min) results in the selective shutdown of canonical mTORC1 signaling and protein translation (Figures 4 and 5) to lower the proteostatic burden and enable the induction of lysophagy (Figure 6). Damaged lysosomal membranes are finally repaired and/or removed by lysophagy as overall PI(3)P levels resurge within hours after the induction of LMP.

**Figure 7.**
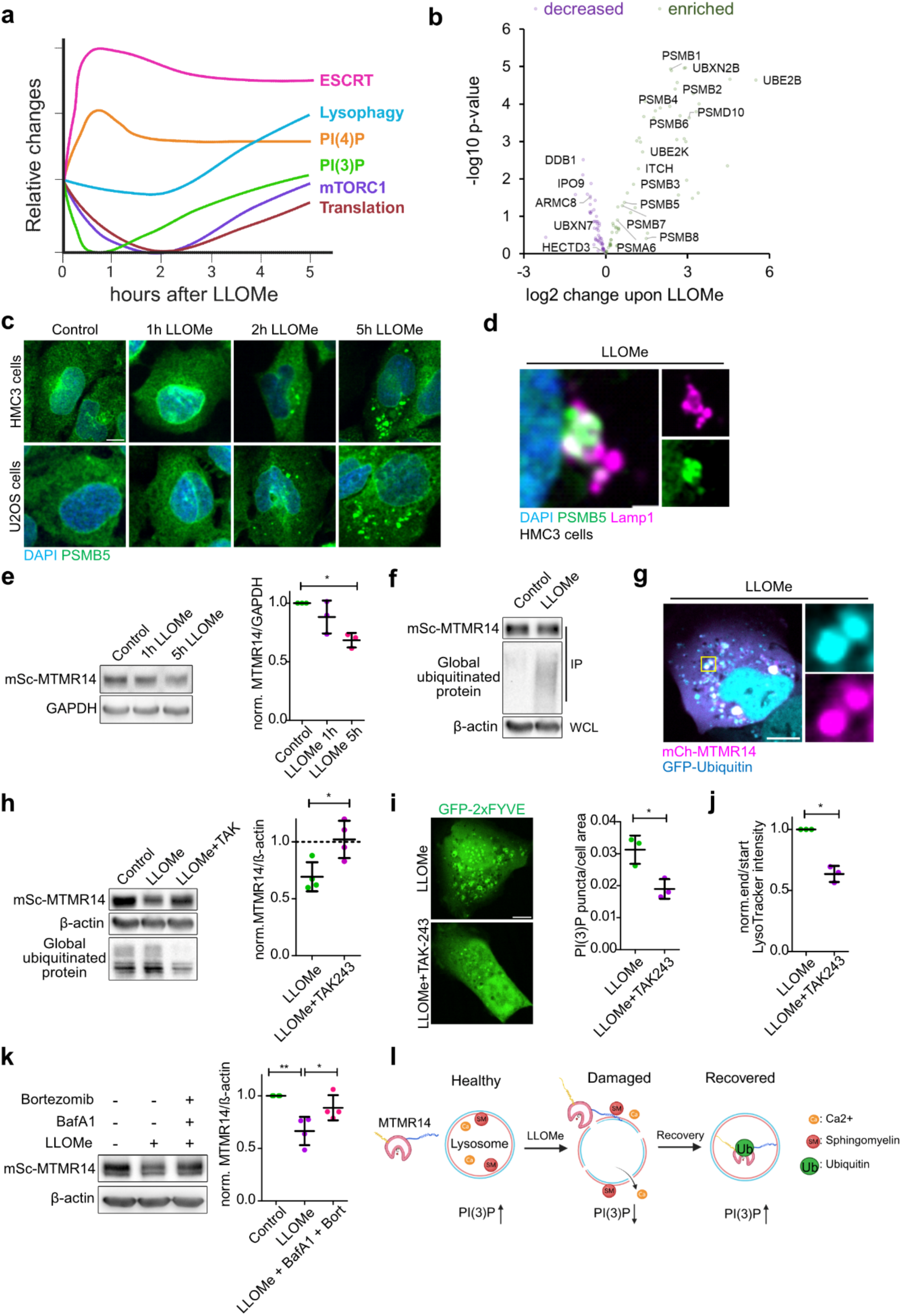
MTMR14 is degraded by ubiquitination to enable PI(3)P recovery. (A) Schematic illustrating the time course of lysosomal membrane damage-induced changes in ESCRTs, lysophagy, PI(4)P, PI(3)P, canonical mTORC1 signalling, and protein translation. Curves were modelled according to experimental data shown in Figure 1c [PI(3)P], Figure 4h (mTORC1), and Figure 6a (Lysophagy), Extended Data Fig. 4h [PI(4)P], Figure 5a (translation), Extended Data Fig. 5a (ESCRT). PI(3)P, mTORC1, and translations curves were normalized between 0 and 1 to allow for comparison of the kinetics. The ESCRT curve represents an average of CHMP4B, IST1, and Alix data (Extended Data Fig. 5a). (b) Volcano plot illustrating the enrichment of proteasome subunits and ubiquitylation machinery on lysosomes immunoprecipitated from control or 1 h 1mM LLOMe-treated HEK293-TMEM192-3xHA cells determined by quantitative proteomics. (c) Representative confocal images of HMC3 and U2OS cells co-stained with DAPI and PSMB5 at indicated time points after 1mM LLOMe treatment. Scale bar, 10 µm. (d) Representative confocal images of 3 h 1mM LLOMe-treated HMC3 cells co-stained for the proteasome subunit PSBM5, Lamp1, and DAPI. Scale bar, 2 µm. (e) Left: representative immunoblot of U2OS-mScarlet-MTMR14 KI cells treated with DMSO control, 1 h 1mM LLOMe, or 5 h 1mM LLOMe. Right: densitometric quantification of mScarlet-MTMR14 relative to GAPDH in control, 1 h and 5 h LLOMe treatment from immunoblots as shown in the left panel. one-way ANOVA (n = 3 independent experiments). (f) Immunoblot of global ubiquitinated protein probed by FK2 antibodies in immunoprecipitation fractions collected from control or 2 h 1mM LLOMe-treated U2OS-mScarlet-MTMR14 KI cells with RFP-trap beads to enrich for mScarlet-MTMR14. (g) Representative confocal images of 2 h LLOMe-treated U2OS cells co-expressing mCherry-MTMR14 and eGFP-Ubiqutin. Scale bar, 10 µm. (h) Left: representative immunoblot of U2OS-mScarlet-MTMR14 KI cells treated with control (DMSO) or 5 h LLOMe with or without 3 µM TAK-243 co-incubation. Right: densitometric quantification of mScarlet-MTMR14 intensity/ ß-actin from immunoblots as shown in the left panel. t test (n = 4 independent experiments). Dotted line denotes mScarlet-MTMR14 level in controls set to 1. (i) Left: representative confocal live cell images of U2OS cells expressing GFP-2xFYVE imaged after treatment with 5 h LLOMe with or without 3 µM TAK-243. Scale bar, 10 µm. Right: number of PI(3)P puncta/cell area quantified in cells as shown in the left panel, fold change over control. t test (n = 3 independent experiments, total number of cells is 108 in LLOMe and 115 in LLOMe + TAK-243 conditions. (j) Quantification of mean LysoTracker intensity/field of view in U2OS cells treated with LLOMe for 5 h with or without 3 µM TAK-243 co-incubation, fold change over control. t test (n = 3 independent experiments, each data point represents 30 fields of view, one field of view containing 35-50 cells, with a size of 3328 × 3328 µm). (k) Left: representative immunoblot of U2OS-mScarlet-MTMR14 KI cells treated with DMSO control, 5 h LLOMe, or 5 h LLOMe with 200 nM Baf A1 and 2 µM Bortezomib. Right: densitometric quantification of mScarlet-MTMR14 relative to GAPDH from immunoblots as shown on the left panel. one-way ANOVA (n = 3 independent experiments). (l) Schematic illustration of MTMR14 recruitment to damaged lysosomes via calcium and sphingomyelin leading to local hydrolysis of PI(3)P during early lysosome damage. Upon prolonged lysosome damage, MTMR14 becomes ubiquitinated, and PI(3)P levels are restored. Statistical analyses were performed using GraphPad Prism. Two-tailed unpaired t-test, paired t-test or one-sample t-tests were conducted using column statistics to compare the sample means to a hypothetical value of 1 or one-way ANOVA with Tukey’s multiple comparisons test. All bar graphs represent mean ± SD unless otherwise stated. ***p < 0.001, **p < 0.01, *p < 0.05. See also Extended Data Fig. 9.

Based on these data, we hypothesized that MTMR14 must be inactivated or removed during the progression of the lysosomal damage response of cells, for example by ubiquitin-mediated protein turnover, i.e. a degradative process that may partially compensate for impaired lysosomal degradation of macromolecules in cells subjected to LMP (in agreement with ^28, 64^). To test this, we conducted unbiased proteomic analysis from cells subjected to damage. This analysis revealed the abundant presence of proteasome subunits as well as ubiquitin ligases and their binding partners on lysosomes (Figure 7b; Extended Data Fig. 9a). Consistently, we found the proteasome subunit PSMB5 to be recruited to lysosomes in both HMC3 or U2OS cells subjected to sustained damage over 5 h (Figure 7c,d), while total PSMB5 expression levels remained unaltered (Extended Data Fig. 9b). These data suggest that local turnover of proteins such as MTMR14 by the ubiquitin-proteasome system may contribute to the response of cells suffering from sustained lysosomal membrane damage. To test directly whether MTMR14 levels are altered during sustained lysosomal membrane damage, we used knock-in cell lines endogenously expressing mScarlet-MTMR14. We found that induction of lysosomal membrane damage resulted in the gradual decline of endogenous mScarlet-MTMR14 levels over hours (Figure 7e) and this was paralleled by LLOMe-induced mScarlet-MTMR14 ubiquitylation (Figure 7f). Moreover, mCherry-MTMR14 colocalized with lysosomal eGFP-ubiquitin puncta in LLOMe-treated cells (Figure 7g). These data suggest that MTMR14 may be subject to targeted degradation by ubiquitin-mediated protein turnover. Indeed, blockade of protein ubiquitination by the E1 ubiquitin-activating enzyme inhibitor TAK-243 occluded the LLOMe-induced degradation of MTMR14 (Figure 7h) and prevented the recovery of endo- and lysosomal PI(3)P levels in cells 5 h post-damage induction by LLOMe (Figure 7i). Importantly, inhibiting protein ubiquitylation also impaired recovery of lysosome function monitored by lysotracker (Figure 7j). Finally, we found that while inhibition of either proteasomal (i.e via Bortezomib) or lysosomal degradation (i.e. via Bafilomycin A1) pathways was insufficient to block LLOMe-induced MTMR14 turnover (Extended Data Fig. 9c,d), combined blockade of both ubiquitin-proteasome and lysosomal proteolysis partially occluded the degradation of MTMR14 in cells subjected to long-term membrane damage (Figure 7k).

Collectively, these results suggest a model (Figure 7l) according to which MTMR14 is rapidly recruited to damaged lysosomes via direct calcium-dependent association with cytosolically exposed sphingomyelin. As damaged lysosomes are repaired or removed, MTMR14 undergoes local ubiquitin-mediated turnover via lysosomal recruitment of proteasomes and the regain of lysosomal proteolysis to facilitate recovery of lysosomal PI(3)P levels, mTORC1 reactivation, and the restoration of lysosome function.

## Discussion

Our study identifies MTMR14, a phosphoinositide 3-phosphatase implicated in human diseases including hereditary myopathy and cancer^46, 53-55^, as part of a damage-induced phosphoinositide module that contributes to how cells cope with lysosomal membrane permeabilization. Specifically, we show that: (i) MTMR14 is rapidly recruited to damaged lysosomes through a coincidence-detection mechanism involving calcium leakage and exposure of sphingomyelin on the cytosolic leaflet, where it promotes local PI(3)P hydrolysis and favours PI(4)P accumulation via PI4K2A ^47^. This MTMR14-driven PI(3)P-to-PI(4)P switch is required for PI4P-dependent ER–lysosome contacts and lipid-transfer–based repair, while ESCRT recruitment proceeds largely independently of MTMR14. (ii) MTMR14-mediated PI(3)P depletion attenuates canonical mTORC1–S6K signalling and global protein synthesis, enabling proteostatic adaptation, lysophagy induction, and improved cell survival after lysosomal membrane permeabilization. (iii) Moreover, we find that prolonged damage triggers ubiquitylation and degradation of MTMR14 at lysosomes, permitting recovery of PI3P and mTORC1 signalling and closing a damage-responsive phosphoinositide module distinct from the nutrient-controlled MTMR14 circuit described previously ^47^. (iv) Finally, our data support a pathophysiological role of the MTMR14-regulated signaling lipid network in the modulation of the cellular response to bacterial pathogens as exemplified here for *Salmonella enterica* ^65-67^. These findings position MTMR14-driven PI(3)P turnover as a local phosphoinositide mechanism that couples lysosomal membrane repair to proteostatic adaptation in response to lysosomal membrane damage.

MTMR14 recruitment relies on signals that have been implicated as very early indicators of lysosomal injury ^23, 68-70^. Previous work has shown that LMP causes rapid calcium leakage and sphingomyelin scrambling at lysosomes, and that Sphingomyelin exposure can help recruit repair factors ^23, 68-70^. Consistent with this, we find that MTMR14 directly binds Sphingomyelin-containing membranes in a calcium-dependent fashion *in vitro*, and that genetic depletion of Sphingomyelin almost completely abolishes MTMR14 recruitment to damaged lysosomes in cells (Figure 2). While this establishes a plausible mechanism for local targeting, we cannot exclude additional determinants, such as other lipids or protein cofactors, especially in tissues and cell types not examined here. Our previous work demonstrated mild starvation-induced MTMR14 recruitment to lysosomes dependent on a preexisting pool of PI(4)P^47^ suggesting that MTMR14 is nested into a feedforward loop that amplifies lysosomal PI(4)P through inhibition of the mTORC1-S6K-PI4K2A axis. Together, these findings argue for a model in which MTMR14 is both upstream and downstream of PI(4)P.

Our data link MTMR14-dependent PI(3)P turnover to several established components of the lysosomal damage response, but also clarify that MTMR14 is unlikely to act alone. Repair via ESCRT ^18, 19^ and the PITT pathway ^21, 22^, CASM ^24, 25^ and stress-granule–mediated stabilization ^10, 26^, lysophagy, and Galectin-^15, 34^ and TFEB-dependent signalling ^16^ all contribute to LMP adaptation. We find that MTMR14 loss markedly blunts damage-induced PI(4)P accumulation and ER–lysosome contacts, yet does not prevent ESCRT recruitment (see Extended Data Fig. 5), indicating that MTMR14 feeds into the PI(4)P-dependent repair arm ^21, 22^. Likewise, MTMR14-dependent repression of canonical mTORC1–S6K signalling co-exists with other modes of mTORC1 regulation on damaged lysosomes, including Galectin-8–GALTOR–mediated inhibition ^13, 34, 35^ and non-canonical TFEB-dependent transcriptional programs ^15, 16^. In line with this, our data show that MTMR14 downtunes canonical mTORC1–S6K signalling in response to LMP. LLOMe-induced lysosomal damage rapidly reduces phosphorylation of S6K and ribosomal S6, and this repression is largely lost when MTMR14 is deleted, but can be phenocopied by genetic or pharmacological manipulations that prevent mTORC1 inhibition (such as TSC2 loss) and rescued by mTORC1/S6K inhibitors (see Figure 4). These findings argue that MTMR14 contributes to canonical mTORC1 inactivation at damaged lysosomes, likely by lowering local PI(3)P levels (in line with ^47^), but they do not exclude additional inputs, including changes in amino-acid efflux, Rag GTPase signalling, Galectin-8–GALTOR ^34^, or lysosome positioning ^31, 32^. mTORC1–TFEB signalling ^71, 72^, in contrast, is largely insensitive to MTMR14 loss and to acute PI(3)P inhibition in our assays, underscoring that distinct phosphoinositide inputs and effector pathways shape canonical versus non-canonical mTORC1 outputs during the damage response. Further investigation is needed to determine how TFEB-mediated lysosome biogenesis^6, 73^ and autophagic lysosome reformation^16^ are temporally and spatially coordinated during the membrane damage response.

A central functional consequence of MTMR14-dependent mTORC1 repression in our system is the rapid inhibition of mTORC1-regulated protein synthesis. We observed a strong, MTMR14-dependent reduction in puromycin incorporation upon LMP, and genetic disruption of downstream translational brakes (4E-BP1/2) or pharmacological reversal of stress-induced translation repression (ISRIB) both compromised lysosome recovery and increased cell death. These data (Figure 5) are consistent with the idea that an acute, mTORC1-linked reduction in global protein synthesis can be protective when lysosomal degradative capacity is impaired, by lowering proteostatic load while repair and removal pathways operate ^37-39^. Future experiments will be needed to examine how selective mRNA translation is remodelled, whether similar translational control occurs *in vivo* under physiological or disease-related lysosomal stress, and which of the four integrated stress response kinases (GCN2, PERK, PKR, and HRI) are involved in the response to LMP.

Intriguingly, we find that damaged lysosomes accumulate ubiquitin ligases, ubiquitin conjugates and proteasome subunits (Figure 7), and that MTMR14 itself becomes ubiquitylated and degraded over time through combined proteasomal and lysosomal pathways. Blocking ubiquitin activation prevents MTMR14 turnover, interferes with recovery of PI(3)P levels, and impairs restoration of lysosomal function after prolonged damage. These observations suggest that ubiquitin-mediated clearance of MTMR14 is one element of a broader proteostatic programme at damaged lysosomes (in line with ^12, 15, 27, 64, 68^), allowing PI(3)P levels and mTORC1 signalling to recover once repair and lysophagy have progressed. Identifying the responsible E3 ligases or the full spectrum of lysosomal and cytosolic substrates that are turned over in this context, as well as disentangling the relative contributions of local proteasomal activity versus global proteasome redistribution during LMP will be exciting avenues for future research.

Finally, our data raise the broader possibility that similar lipid-driven modules operate at other damaged membranes. We find that MTMR14 can be recruited to permeabilized plasma membranes under conditions that promote calcium influx and Sphingomyelin exposure, suggesting that the same coincidence logic may apply to non-lysosomal compartments. Whether MTMR14-dependent PI(3)P turnover contributes to plasma membrane repair, organelle crosstalk, or proteostasis in tissues subject to mechanical or proteotoxic stress—such as skeletal muscle, microglia in the brain, or tumour microenvironments—remains an open question. Intriguingly, recent work identified LMP as a targetable pathway in hereditary muscular dystrophies ^5^ suggesting that myopathies caused by MTMR14 loss-of-function ^54^ could benefit from targeting mTORC1-S6K signaling. Addressing this will require *in vivo* imaging and genetic perturbation of MTMR14 under physiological damage regimes, including contraction-induced injury, aggregation-prone protein accumulation and infection. Taken together, our work defines MTMR14-dependent PI(3)P turnover on damaged lysosomes as a local phosphoinositide module that promotes lysosomal membrane integrity and couples canonical mTORC1 signalling to proteostatic adaptation, while leaving room for additional pathways and cell-type-specific mechanisms that remain to be elucidated.

## Methods

### Materials

#### Plasmids

YFP-MTMR14WT and YFP-MTMR14(C330S) plasimds as a gift from Jocelyn Laporte (IGBMC Strasbourg, France), eGFP-EqtSM ^23^, eGFP-2xFYVE (in-house), pmCherry-Gal3 (addgene #85662), eGFP-OSBP-PH (in-house), GFP-SNXA ^74^ was a gift from Jason S. King (University of Sheffield, UK), eGFP-KIF16Btail (addgene #134624), mCherry-MTMR14 (in-house), pmCherry-P4M-SidMx2 (in-house), pRK5-eGFP-Tau P301L (addgene # 46908), eGFP-MTM1 (in-house), eGFP-MTMR1 (in-house), eGFP-MTMR2 (in-house), eGFP-MTMR6 (in-house), EGFP-MTMR7 (in-house), eGFP-MTMR8 (in-house), pMRX-IB-TMEM192-mKeima (addgene #215359), HA-PI4K2A (in-house), eGFP-Ubiquitin (addgene #187910), mCherry-MTMR14 ΔN (delete 1-68) (in-house), mCherry-MTMR14 ΔC (delete 406-650) (in-house), mCh-MTMR14 Δ620-650 (in-house), mCh-MTMR14 Δ586-650 (in-house), CFP-MTMR14 C-terminus (delete 1-570) (in-house), mCh-MTMR14 C-terminus (delete 1-570) (in-house), mCh-MTMR14 C-terminus 7xmut (L590E L591E A592S A593S L594E F608E Y612E) (in-house), mCh-MTMR14 FL 7xmut (L590E L591E A592S A593S L594E F608E Y612E) (in-house). All constructs were verified by dsDNA sequencing.

#### Antibodies

GAPDH (glyceraldehyde-3-phosphate dehydrogenase; mouse, Sigma-Aldrich, G8795, 1:4000); actin ( mouse, Sigma-Aldrich, Cat# A5441, 1:4000);Phospho-p70 S6 Kinase (Thr389) (rabbit, Cell signaling, Cat# 9205L,1:500), p70 S6 Kinase Antibody (rabbit, Cell Signaling, 9202L, 1:1000); PI4K2A (homemade); MTMR14 (homemade); dsRed/mScarlet (rabbit, Clontech, Cat# 632496, 1:1000); pULK(S757) (rabbit, Cell signaling, Cat# 6888, 1:1000); ULK1(rabbit, Cell signaling, Cat# 8054, 1:1000); p4E-BP1(S65) (rabbit, Cell signaling, Cat# 13443; 1:500); 4EBP1 (rabbit, Cell signaling, Cat# 9644; 1:500); pTFEB(S211) (rabbit, Cell signaling, Cat# 37681, 1:1000); TFEB (rabbit, Biomol, Cat# A303-673A, 1:1000); Anti-mouse HRP (Jackson, Cat# 115-035-003, 1:4000); Anti-rabbit HRP (Jackson, Cat# 111-035-003, 1:4000); Anti-mouse Alexa568 (IgM-specific) (Thermo, Cat# A21043, immunofluorescence 1:400); PtdIns(4)P (Echelon, Cat# z-p004, immunofluorescence 1:200); a-RXRXX[S/T] (Cell Signaling, Cat# 10001, 1:500); pS6 (Cell Signaling, Cat# 2215, 1:1000); pS6(S240/244) (Cell Signaling, Cat# 5364, 1:1000); puromycin (Kerafast, Cat# Kf-Ab02366-23.0, WB 1:500, immunofluorescence 1:2000); CHMP4B (Proteintech, 13683-1-AP, immunofluorescence 1:200); IST1 (proteintech, Cat# 19842-1-AP, immunofluorescence 1:200); Alix (Biolegend, Cat# 634501); PSMB5 (Thermo, Cat# PA1-977, immunofluorescence 1:200); Anti-mouse CF568 (Biotium, Cat# 20109-0.25ml, immunofluorescence 1:1000); Anti-rabbit Alexa 647 (Thermo, Cat# A31573, immunofluorescence 1:1000); Anti-rabbit Alexa 568 (Thermo/ Molecular Probes, Cat# A11036, immunofluorescence 1:1000); Anti-mouse Alexa 647 (life, Cat# A21238, immunofluorescence 1:1000); anti-Mouse IRDye 680 CW (LICOR, P/N 925-68070, 1:4000); anti-Rabbit IRDye 680 CW (LICOR, P/N 925-68180, 1:4000).

#### Chemicals

L-Leucyl-L-Leucine methyl ester hydrobromide (LLOME, Sigma, Cat# L7393, 1mM, 2mM), Gly-Phe-β-naphthylamide (GPN, Cayman Chemical, Cat# 14634, 200µM), O-methyl-serine dodecylamide hydrochloride (MSDH, Avanti Polar Lipids, Cat# 850546, 50µM), Benzalkonium chloride (BAC, CAS# 63449-41-2, MP Biomedicals, Santa Ana, CA, USA, 5ug/mL), hydrogen peroxide (H_2_O_2_, Cat# ‘9681.2, Roth, 0.5mM), VPS34-IN1 (Selleckchem, Cat# S7980, 5µM), apilimod (Echelon Biosciences, Cat# B-0308, 50nM), rapamycin (Santa Cruz Biotechnology, Cat# sc-3504, 100nM), AZD8055 (AdooQ Bioscience/hölzel, Cat# A10114, 200nM), Anisomycin (Sigma, Cat# A9789, 30µM), LY2584702 (S6Ki, Selleckchem, Cat# S7704, 2µM, 5µM), MRT68921 dihydrochloride (MedChemExpress, Cat# 2080306-21-2, 1µM), bortezomib (MedChemExpress, Cat# 179324-69-7, 2µM), CCCP (Sigma, Cat# C2759-100MG, 10µM), Bafilomycin A1(Sigma, Cat# SML1661, 100nM or 200nM), nigericin (Invivogen, Cat# tlrl-nig,5µM), D-Mannitol (Sigma, Cat# M4125, 0.25M), Puromycin (InvivoGen, Cat# ant-pr-1, 2µM), Ionomycin (Calbiochem, Cat# 407952, 5µM), Doxycycline (Sigma, Cat# D-9891, 1µg/mL), Digitonin 5%(Thermo, Cat# BN20061, 20 mM). BAPTA-AM (Cayman Chemical, Cat# 15551, 100µM), MLi-2 (MedChemExpress, Cat# 1627091-47-7, 5µM). Chemicals were directly added to cell culture media to induce lysosomal membrane damage with equal volume of vehicle (DMSO or water) added into control wells.

#### siRNA

siRNA used were: Scrambled siControl 5’-CGUACGCGGAAUACUUCGA-3’, siMTMR14 5’-GGACAAAGCCCAGCGCUAU-3’, siMTMR14 (Horizon Dharmacon, Cat# L-016118-00-0010) and siTFEB (Horizon Dharmacon, Cat# L-009798-00-0010) in U2OS cells, 2.5 × 10⁵ to 3 × 10⁵ cells were seeded per well in 6-well plates and reverse transfected with 50nM siRNA using 6 µl INTERFERin pre-diluted in serum-free medium. A forward transfection was performed on day two, and cells were analyzed on day four. For transient knockdown of MTMR14 (Horizon Dharmacon, Cat# L-050513-01-0010) and CHMP4B (Horizon Dharmacon, Cat# L-041531-01-0010) in C2C12 cells, siRNA transfection was performed using Lipofectamine RNAiMAX (Thermo Fisher Scientific) according to the manufacturer’s instructions.

#### Cell lines

Cell lines U2OS, HeLa, APRE-19, C2C12, BV2 and HMC3 cells were obtained from ATCC. HeLa SMS1/SMS2 double-KO (SMS-DKO) and U2OS SMS1/SMS2 double-KO (SMS-DKO) cells were previously described^23^. HeLa TSC2 KO cells were a gift from Elizabeth Henske (Harvard Medical school, USA). The cell lines in this study have different morphologies and growth rates, and contamination was consistently monitored. All cell lines used in this study were confirmed to be free of mycoplasma contamination by PCR detection and were routinely maintained using a mycoplasma prevention reagent. U2OS, HeLa, APRE-19, C2C12, BV2 and HMC3 were maintained in DMEM containing 4.5 g/L glucose and L-glutamine (Gibco), supplemented with 10% heat-inactivated fetal calf serum (Gibco), 100 U/mL penicillin, and 100 µg/mL streptomycin (Gibco). All cells were maintained at 37 °C with 5% CO_2_.

#### Bacterial strains

*Salmonella enterica* Typhimurium strains *ΔsifA*, a kind gift from Dirk Bumann (Biozentrum, University of Basel, Switzerland).

#### Software used

Fiji (https://imagej.net/downloads), LOG 3D Fiji plugin (http://bigwww.epfl.ch/sage/soft/LoG3D/), JaCoP Fiji Plugin, Affinity, Biorender.

## Methods

### U2OS knockout cell line generation

#### Guide DNA design

The single guide RNAs (sgRNA) were designed with the online tools ChopChop v3 using options for KI resp. KO (http://chopchop.cbu.uib.no/; Labun et al., 2019) and CRISPOR (http://crispor.tefor.net/; Concordet, J.-P.,Haeussler, M., 2018) (Sequences for sgRNAs are listed in supplementary table 3.

To generate sgRNA expression plasmids (=Guide), oligonucleotides containing sgRNA sequence were cloned into a pSpCas9(BB)-2A-GFP vector (Addgene plasmid #48138; a gift from Feng Zhang) using *BbsI* restriction digest. The GFP tag of the guide vector was used for single cell sorting.

#### Cell culture and transfections

U2OS cells were maintained in Dulbecco’s Modified Eagle’s Medium (DMEM, high glucose) (Gibco) supplemented with 10% fetal bovine serum (FBS) (Sigma) and 1% penicillin-streptomycin (Gibco) in a 5% CO_2_ atmosphere at 37 °C. Cells were seeded in 100-well plates (2 million per plate). Transfection was done 1 day later with polyethylenimine (PEI, Polysciences) and the respective DNA. 48 hours after transfection, single cells were sorted into 96-well plates using the FACSMelody (BD Biosciences). After single cells are grown, positive clones are detected by sequencing (MTMR14-KO).

### Transient transfection

Cells were seeded on Matrigel (BD/Corning)-coated 8-well chamber slides (ibidi, 80827). The next day, expression plasmids were transfected into cells using Lipofectamine2000 according to the manufacturer’s instructions.

### Generation of Doxycycline-inducible stable cell lines

Lentivirus was generated by transient transfection of HEK293T cells, seeded in 10-cm cell culture plates at 80–90% confluency, using 3.5 µg of pCMV delta R8.2, 0.5 µg of VSV-G, and 4 µg of pLVX-TetONE-puro–based constructs. Plasmids were mixed with 16 µl of JetPrime in 400 µl of JetPrime buffer. After 16 hours of transfection, the medium was replaced with 7 ml of fresh DMEM. After 48 hours, the viral supernatant was collected, and cellular debris was removed by centrifugation at 3,000 rpm for 20 minutes. Approximately 4–5 ml of the clarified viral supernatant was added to target cells in the presence of polybrene (Merck, Cat. #TR-1003-G) at a final concentration of 10 µg/ml. Cells were incubated with the virus for 16 hours and then replenished with fresh DMEM containing puromycin (2 µg/ml) for selection over 2-3 days. Following selection, cells were maintained in culture without puromycin for an additional 3–4 days to allow stabilization. For induction of protein expression, doxycycline (1 µg/ml) was added for 16 hours.

### Light microscopy

#### Immunocytochemistry

Cells were seeded on Matrigel (BD/Corning)-coated 12mm coverslips in 24 well dishes, fixed in 4%paraformaldehyde/4% sucrose in PBS for 15 minutes at room temperature, washed 3 times with PBS, incubated 1h room temperature with primary antibodies (1:200 in 10% goat serum/0.05% Saponin in PBS), washed 3 times 5 minutes with PBS, incubated 1h room temperature with fluorophore-conjugated secondary antibody 1:1000 in 10% goat serum/0.05% Saponin/PBS including DAPI, washed 3 times 5 minutes with PBS, and mounted with Shandon Immu-Mount^TM^.

#### PI(4)P and LAMP2 co-staining and GFP-2xFYVE overlay

We essentially followed a previously published protocol^47^. Briefly, cells were grown on 12 mm glass coverslips in 24-well plates. After a single PBS wash, cells were fixed for 15 minutes at room temperature in 2% paraformaldehyde (PFA) and 2% sucrose in 1× PBS. Following fixation, cells were washed three times with 50 mM NH₄Cl in 1× PBS to quench residual PFA. Cells were then permeabilized for 5 minutes at room temperature with 20 mM digitonin (prepared from a 5% stock; Invitrogen) diluted in Buffer A (20 mM PIPES, pH 6.8 [adjusted with NaOH], 137 mM NaCl, 2.7 mM KCl). After permeabilization, cells were washed three times with Buffer A. Blocking was performed for 1 hour at room temperature using 5% goat serum in Buffer A with 0.25mg/ml of purified GFP-2xFYVE (Hrs). Cells were then washed twice with Buffer A. Primary antibodies against PI(4)P and LAMP2 were diluted in 5% goat serum in Buffer A and incubated with the cells for 1 hour at room temperature. After three washes with Buffer A, cells were incubated for 1 hour at room temperature with fluorophore-conjugated secondary antibodies diluted in 5% goat serum in PBS containing DAPI. Following three washes with PBS, cells were post-fixed in 2% PFA and 2% sucrose in 1× PBS, then washed again with 50 mM NH₄Cl in 1× PBS. Coverslips were mounted using Shandon™ ImmuMount™ mounting medium.

#### TMEM192-mKeima live imaging

TMEM192-mKeima expressing cells were precultured overnight in the aforementioned DMEM on Matrigel-coated 8-well chamber slides (ibidi, 80827). Cells were treated with 1mM LLOMe to induce lysophagy, followed by culture in DMEM/FBS. After incubation, the cells were subjected to Nikon-CSU Yokogawa Spinning disk CSU-X1 microscope. The cells were examined using using a 40x air objective and hardware autofocus at room temperature. The cells were observed under neutral and acidic conditions using excitation wavelengths of 405 nm and 561 nm, respectively, both with a 600/50 emission filter.

### Live cell imaging

Cells were cultured in seeded on Matrigel-coated 8-well chamber slides (ibidi, 80827). The next day, live-cell imaging was carried out using Nikon-CSU Yokogawa Spinning disk CSU-X1 microscope equipped with an EMCCD Camera (Andor AU-888) or CSU-W1 microscope equipped with 2 backilluminated sCMOS Cameras at 37°C in the presence of 5% CO_2_.

### Lysosome damage and cytotoxicity assays

#### LLOMe washout assay

Cells were treated with 2 mM LLOMe for 30 minutes, followed by five washes with DMEM containing FBS, and then maintained in fresh DMEM/FBS for subsequent analysis.

#### LLOMe long-term treatment assay

Cells were incubated with 1 mM LLOMe in DMEM/FBS for 5 hours.

#### GPN induce lysosome damage model

Cells were treated with 200 nM GPN for the desired time.

#### LDH LLOMe assay

Cells were treated with 4 mM LLOMe in DMEM/FBS for 5 hours to assess lactate dehydrogenase (LDH) release as a marker of cell damage.

#### Overexpression of pRK5-eGFP-tau P301L

tau P301L overexpression was used to induce lysosomal damage. Cells were seeded at approximately 60-70% confluency in 8-well chamber slides (ibidi, 80827). After 18-24 hours, cells were co-transfected with pRK5-eGFP-tau P301L and mCherry-Galectin-3 plasmids using Lipofectamine 2000 (or another transfection reagent) following the manufacturer’s protocol. At 24 or 48 hours post-transfection, lysosomal integrity was assessed by monitoring Galectin-3 (Gal3) recruitment, a marker of lysosomal membrane damage.

### Plasma membrane damage assays

*MSDH induce Plasma membrane model:* Cells were treated with 50µM MSDH for the desired time. MSDH concentrations above 20 µM cause a loss of plasma membrane integrity and are more likely to kill the cell by rupturing the plasma membrane than by entering the cell and inducing LMP through lysosomal accumulation.^75^ *BAC induce Plasma membrane model:* Cells were treated with 5ug/mL benzalkonium chloride (BAC) ^76^ for the desired time.

### Lysotracker recovery assay

Cells were incubated with Lysotracker Red DND-99 (Thermo Fisher, L7528) (1:1000) and Hoechst 33342 (1:200) for 30 minutes. Then, cells were washed with DMEM containing FBS and media were replaced with fresh DMEM containing LysoTracker Red (1:4000).

### Electron Microscopy

Cells were fixed with 2% glutaraldehyde in PBS, postfixed in 1% OsO4 and 1,5% potassium hexacyanoferrate (III) in 0,1M cacodylate buffer on ice for 45 minutes, washed in water, treated with aqueous 1% OsO4 for 20 minutes, stained en bloc with 1% aqueous uranyl acetate, dehydrated in acetone, infiltrated with Durcupan resin. Following resin polymerization, coverslips were removed with help of liquid nitrogene, ultrathin sections were mounted on silicium wafers. Imaging of cells was performed at Helios 5CX scanning microscope using DBS detector. Quantification of ER/lysosome contacts per cell was manually performed with identity of cell being blinded for investigator.

### Protein purification

MTMR14 constructs were expressed in Expi293F cells (Thermo Fisher A14527) by transient transfection. Cells were grown in Expi293 media (Thermo Fisher A1435102) in a Multitron Pro shaker set at 37 °C, 8% CO2 and 110 rpm. Transfections were performed by incubating PEI and DNA constructs for 20 minutes (Polyethyleneimine "MAX", MW 40,000, Polysciences, 24765, total 3 mg PEI/L cells to a ratio of 1 mg DNA/L cells) before adding to EXPI cells at a density of 2.5x106 cells/mL. After 24-48 hours, cells were collected by centrifugation, washed gently with ice cold 1x PBS, and then frozen in liquid N2 and stored at -80 °C.

Frozen pellets were re-suspended in lysis buffer (10 mM HEPES pH 7.5, 150 mM NaCl, 0.4% CHAPS, 0.5 mM TCEP, protease inhibitor (Protease Inhibitor Cocktail Set III, Sigma)) and incubated with rotation for 30 mins at 4C to allow for cell lysis. Cell lysate was centrifuged at 23500 rpm at 4 C for 60 minutes (Thermo Fisher, SA-300 rotor). The supernatant was then loaded onto a 5 mL StrepTrap HP column (Cytiva #28-9075-48) that had been equilibrated in Buffer A (10 mM HEPES pH 7.5, 150 mM NaCl, 0.5 mM TCEP). The column was washed with 50 mL of Buffer A (10 mM HEPES pH 7.5, 150 mM NaCl, 0.5 mM TCEP), before being eluted with Buffer A containing 10 mM d-desthiobiotin. The elution fractions were pooled and concentrated in a 50,000 MWCO Amicon concentrator (Merck Millipore). Protein was aliquoted, snap-frozen in liquid nitrogen, and stored at -80°C or used fresh.

### Protein binding to Small Unilamellar Vesicles (SUVs)

SUVs were prepared from synthetic lipid mixtures containing either 60 mol% dioleoyl-phosphatidylcholine (DOPC; Avanti Polar Lipids, #850375), 20 mol% cholesterol (Avanti Polar Lipids, #700000), and 20 mol% porcine brain sphingomyelin (SM; Avanti Polar Lipids, #860062), or 80 mol% DOPC and 20 mol% cholesterol. Protein buffer: 10mM Hepes pH 7.5, 150mM NaCl, 0.5mM EGTA (Roth). Lipids were hydrated in liposome binding buffer containing 10mM Hepes pH 7.5, 150mM KCl, 0.5mM EGTA. SUVs were prepared by size exclusion chromatography as described^77^.

For binding assays, 60 µL of liposomes (stock lipid concentration: 0.7 mM) were incubated with 4 µg of protein per condition in the presence of 1 mM CaCl₂ at room temperature for 5 min, ultracentrifuge at 70,000rpm, at 4°C for 15 min.

### Cell lysates and immunoblotting

Cells were washed with 2 times cold PBS and lysed in 1% Igepal (Sigma-Aldrich CA-630, formerly known as NP-40) in 50 mM Tris buffer (pH 8.0) containing 150 mM NaCl, supplemented with Roche cComplete protease inhibitors and Sigma-Aldrich phosphatase inhibitor cocktails #2 (P5726) and #3 (P0044). Cell lysates were cleared by centrifugation, and the resulting supernatants were boiled with sample buffer for 10 minutes. Proteins were resolved on 8–12% SDS-PAGE gels and transferred onto nitrocellulose membranes using transfer buffer containing 20% methanol. Membranes were blocked for 1 hour at room temperature in 5% skimmed milk prepared in Tris-buffered saline with 0.1% Tween-20 (TBST). Membranes were then incubated overnight at 4 °C with primary antibodies diluted in blocking buffer. After washing in TBST, membranes were incubated for 1 hour at room temperature with horseradish peroxidase (HRP)-conjugated secondary antibodies. Signals were imaged using a Bio-Rad ChemiDoc system and quantified using Fiji’s Gel Analyzer densitometry tool.

### Lysosome-IP and proteomics

#### Lyso-IP

HEK293 cells stably expressing TMEM192-3×HA were cultured in 15 cm dishes until confluent. Cells were washed twice with cold PBS and collected by scraping into 950 µL of cold KPBS. After centrifugation at 1,000 × g for 3 min at 4 °C, cell pellets were resuspended in 950 µL cold KPBS. A 25 µL aliquot was reserved for whole-cell lysate (WCL) preparation. The remaining suspension was homogenized on ice using a pre-chilled Dounce homogenizer (25 strokes), and the homogenate was centrifuged at 1,000 × g for 3 min at 4 °C. The supernatant containing organelles was incubated with pre-washed anti-HA magnetic beads for lysosome isolation. Beads were separated on a magnetic rack and washed three times with cold KPBS. For improved purity, beads were transferred to a clean tube after the second wash. Bound proteins were eluted in 1% NP-40 buffer supplemented with protease (Roche cComplete) and phosphatase inhibitors (Sigma, Cat. #P5726 and #P0044), along with 2× SDS loading dye. Samples were heated at 95 °C for 10 min and stored at –20 °C until analysis. For WCL preparation, the 25 µL reserved aliquot was lysed in 140 µL of lysis buffer, centrifuged at ∼20,000 × g for 15 min at 4 °C, and the supernatant was mixed with 6× SDS loading dye. Samples were boiled at 95 °C for 10 min and stored at –20 °C.

#### Proteolytic digestion via SP3

Lyso-IPs were eluted using 4X SDS-PAGE Loading dye for 10 min at 95 °C and then digested using the SP3 protocol. Proteins were reduced with 5 mM TCEP and alkylated with 40 mM CAA for 1 h at room temperature. Sera-Mag beads were added to the sample, followed by 50% ACN. After a brief incubation the supernatants were removed and the beads were washed twice with ethanol and once with ACN. Proteins were digested in 50 mM TEAB (pH 8.5) with LysC and Trypsin (both in enzyme:protein ratio 1:50 wt:wt) for 16 h at 37 °C. Beads were washed twice with an excess of ACN. Peptides were eluted with 5% DMSO and dried until further use.

#### Liquid chromatography and mass spectrometry

Desalted peptides were resuspended in 1% acetonitrile (ACN) with 0.05% trifluoroacetic acid (TFA) and injected into a Thermo Scientific Vanquish Neo system coupled online to an Orbitrap Astral mass spectrometer (Tune version 1.1). Prior to separation, peptides were trapped on a PepMap C18 trap column (0.075 × 50 mm, 3 μm, 100 Å, Thermo Fisher). Peptide separation was performed using reverse-phase chromatography on an in-house packed C18 column (Poroshell 120 EC-C18, 2.7 μm, Agilent). Elution was carried out over a 40-minute linear gradient of increasing ACN concentration at a flow rate of 300 nL/min. Data were acquired in data-independent acquisition (DIA) mode. MS1 scans were performed in the Orbitrap at a resolution of 240,000 over a scan range of m/z 430–680, with a 40% RF lens setting, 500% AGC target, and a maximum injection time of 5 ms. MS2 scans were acquired in the Astral analyzer using 2 m/z isolation windows with no overlap, automatic window type, placement optimization enabled, normalized collision energy of 25%, scan range of m/z 145–2000, 40% RF lens, 500% AGC target, 5 ms maximum injection time, and a 0.6-second duty cycle.

#### Identification and quantification of proteins from whole-cell lysates

Raw data were analyzed using DIA-NN version 2.0 ^78^. Spectra were searched against an in silico spectral library generated from the human proteome (retrieved on 2024-04-05) and common contaminants. DIA-NN was configured with the following parameters: MS1 tolerance of 10 ppm, MS2 tolerance of 20 ppm, peptide length range of 7 – 30, trypsin digestion allowing a maximum of one missed cleavage. Match-between-runs was enabled. Variable modifications included oxidation of methionine (+15.9949 Da) and protein N-terminal acetylation (+42.0106 Da), while carbamidomethylation of cysteine (+57.0215 Da) was set as a fixed modification. All other settings were left default. Quantitation analysis was performed in R. Proteins were retained if quantified in at least two out of three replicates in at least one condition. Intensity values were log2-transformed, normalized using variance stabilizing normalization (VSN), and missing values were imputed with random draws from a gaussian distribution centered at the minimal value of each sample. Differential expression analysis was conducted using the limma package to compute log2 fold changes and p-values. Gene Ontology (GO) terms were annotated using the org.Hs.eg.db package.

#### Data availability

The mass spectrometry data have been deposited to the ProteomeXchange Consortium via the PRIDE partner repository with the identifier: PXD065904.

### Salmonella assays

#### Preparation of Salmonella

A single colony of *Salmonella enterica* Typhimurium was inoculated into LB medium (Carl Roth, Karlsruhe, Germany) in a 15 mL tube with a loose cap and incubated overnight at 37 °C with shaking. The next day, a portion of the overnight culture was transferred into fresh LB medium (1:33 dilution) and incubated statically at 37 °C for 3 hours. After incubation, bacteria were pelleted by centrifugation at 10,000 × g for 2 minutes, washed once with PBS, and resuspended in cell culture medium. The concentration of bacteria was measured at OD 600nm, considering that an OD 600nm of 1.0 corresponds to approximately 1.2 × 10⁹ CFU/mL.

#### Infection of human cells

U-2OS cells were plated one day prior to infection, and on the day of infection, if the cells are cultured in medium containing antibiotics, cells were washed with antibiotic-free medium. *Salmonella* were diluted in cell culture medium at a multiplicity of infection (MOI) of 10:1 (10 bacteria per 1 human cell), added to cells, and incubated for 30 minutes at 37 °C. Afterwards, to kill extracellular bacteria, cells were washed twice with PBS and cell culture medium containing 50 µg/mL gentamicin (Gibco) was added and incubated for 40 minutes at 37 °C. After this, the medium was replaced with medium containing 5 µg/mL gentamicin and incubation continued for the desired duration at 37 °C.

#### Colony-forming unit (CFU) assay

At the time point of interest, cells were washed twice with PBS and lysed with 1% Triton X-100 and 0.1% SDS in PBS. The lysate was serially diluted in PBS and 10 µL of each dilution were spotted (at least three spots per dilution) onto LB agar plates (Carl Roth). Plates were incubated overnight at 37 °C and the resulting colonies were counted to calculate the colony-forming units (CFU) from both the dilution factor and the spot volume. The CFU counts were normalize to the value at the 1 hour or 1.5h post-infection time point to compute bacterial intracellular growth rates.

### Cytotoxicity assay (LDH release)

Seed human cells and incubate with appropriate stimulus. Use all conditions in triplicates (technical replicates). Include addition of untreated cells and lysed cells, too, in triplicate. When the right incubation time has elapsed, harvest the cell culture supernatant. If cells were infected by S2-grade bacteria, filter the supernatant on 0.22 µm filter. Filtering is unnecessary for S1-grade experiment. For untreated or uninfected cells, take supernatant as a low LDH control. For high LDH control, treat with 2% Triton X-100 in medium, incubate for 5 minutes, and then take the supernatant. Also, take freshly taken cell-treated free medium as a blank (cell-free control). To measure lactate dehydrogenase (LDH) release, apply the Cytotoxicity Detection Kit (LDH) (Roche, Cat. No. 11644793001). Thaw out all kit components and reconstitute Bottle 1 by adding 1 mL of distilled water. Reconstitute the reaction buffer by mixing 2.5 µL of Bottle 1 with 112.5 µL of Bottle 2 per reaction. Mix in a 96-well plate 100 µL each of supernatant with 100 µL of the reaction buffer. Incubate at 37 °C for up to 30 minutes. Read absorbance at 492 nm on a plate reader.

### Gene expression analysis in human muscle samples

Whole-genome RNA sequencing was performed in human muscle samples (m. quadriceps femoris) from a prospective monocentric cohort study as previously described (Baritello et al., JCSM, in revision). In brief, the study included 63 patients (≥ 70 years of age) admitted to the Department of Cardiovascular Surgery, Charité-Universitätsmedizin Berlin, for elective cardiac surgery (e.g. coronary artery bypass grafting) or minimally invasive procedure (i.e. TAVI). The datasets are available from the GEO database under GEO accession number: GSE287726.

### Lipid analysis

To assess sphingomyelin (SM) biosynthesis U2OS SMS-DKO cells were metabolically labeled with a clickable sphingosine analogue (4 uM, 16 h) as in Kol et al. ^79, 80^. Total lipids were extracted, click-reacted with 3-azido-7-hydroxycoumarin^80^, separated by TLC, and analyzed by fluorescence detection.

Sphingomyelin (SM) and hexosylceramide (HexCer) content in total lipid extracts from wildtype (WT) and SMS-DKO cells were determined by liquid chromatography with mass spectrometry (LC-MS/MS) as described^23^ and expressed in pmol per 100 pmol of total phospholipid analyzed.

### Image analysis

#### MTMR14 recruitment

Image analysis was performed using ImageJ. YFP-MTMR14 (or mCherry-MTMR14)-decorated vesicles were filtered and detected using a LoG filter and the Find Maxima command, respectively, and counted before and after LLOMe treatments.

#### PI(3)P level

Image analysis was performed using ImageJ. GFP-2xFYVE-decorated vesicles were filtered and detected using a LoG filter and the Find Maxima command, respectively, and counted before and after LLOMe treatments.

#### PI(4)P staining

Image analysis was performed using ImageJ. Background signal was subtracted, and thresholding was applied to enhance signal detection. A fixed-size region of interest (ROI), usually covering the entire frame, was uniformly applied across image stacks. Mean intensity values within the ROI were measured using the Multi Measure function.

#### Lysotracker recovery assay

Background subtraction was performed on the image stack. Mean intensity measurements within the ROI were obtained using the ROI Manager’s Multi Measure function in ImageJ.

#### Galectin3 puncta, TMEM192-mKeima puncta

Image analysis was performed using ImageJ. mCherry-*Galectin3*-decorated vesicles or *TMEM192-mKeima*-decorated vesicles were filtered and detected using a LoG filter and the Find Maxima command, respectively, and counted before and after LLOMe treatments.

#### Pearson’s correlation

Pearson’s correlation was calculated using the Fiji JaCoP Plugin.

### Additional Software

Cartoons and schematics were generated using BioRender and Affinity.

### Statistical analysis

All data are presented as mean ± SD, sample numbers are indicated in the figure legends. The number of replicates is indicated in the figure legends. Statistical significance was evaluated using Student’s t-test, or one-sample t-tests were conducted using column statistics to compare the sample means to a hypothetical value of 1, or one-way ANOVA with Tukey’s multiple comparisons test (to compare the mean of each column with every other column). All of the statistical tests were calculated in Graph Pad Prism Significant differences were marked as * p<0.05, ** p<0.01, *** p< 0.001.

## Acknowledgments

We thank Drs. Elizabeth P. Henske and Nicola Alesi (Harvard Medical School, Boston, MA) for the kind gift of TSC2 KO HeLa cells, Silke Zillmann, Delia Löwe, Maria Mühlbauer, and Claudia Schmidt for expert technical assistance, and Drs. Martin Lehmann and Tolga Soykan for aid with mircoscopy and image analysis. Y.S. was supported by a scholarship from the Chinese Scholarship Council (CSC). V.H. acknowledges funding by the Deutsche Forschungsgemeinschaft (TRR186/ A08).

## Author contributions

M.E., Y.S., and V.H conceived the project and designed the experiments. Y.S. performed most of the experiments. D.P. performed and analysed together with Y.S. electron microscopy experiments. J.M.V. and I.D., together with Y.S., designed and performed the experiments with bacteria. O.A.A., E.S. and J.C.M.H. generated and characterized SMS1/2 DKO cell lines. M.R. and F.L. performed organelle proteomics together with Y.S.. Y.S., M.E. and V.H. wrote the manuscript and drafted the figures with input from all authors.

## Competing interests

The authors declare no competing financial interests.

**Supplementary Information** is linked to the online-version of the paper.

**Correspondence and requests for materials** to *V.H. or M.E.*.

## Data availability

All data are available in the main text or the supplementary materials. Materials and reagents are available from the corresponding author upon request.

## Extended Data Figures

**Extended Data Fig. 1.**
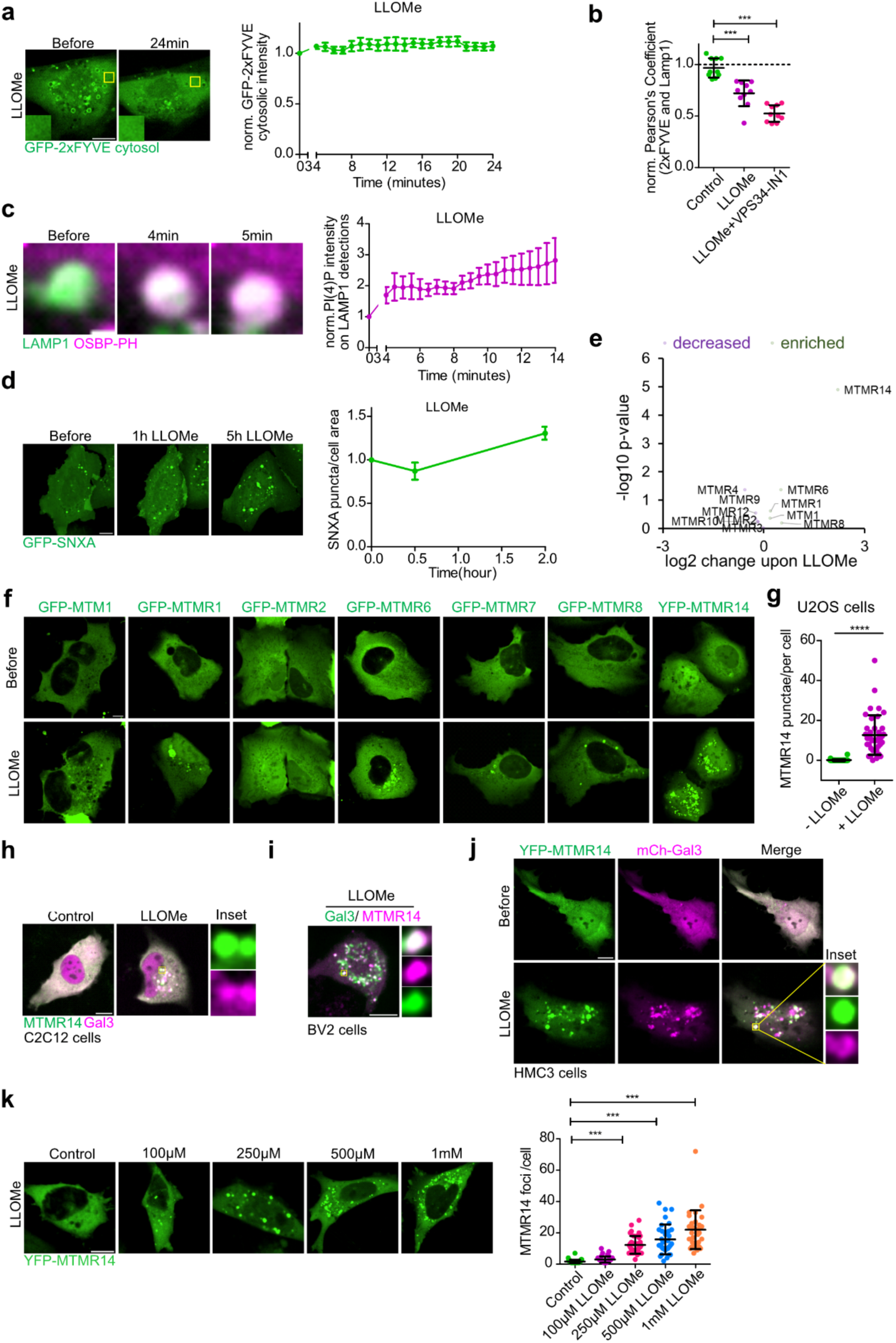
Phosphoinositide and MTMR14 dynamics during the lysosomal damage response. (a) Left: eGFP-2xFYVE cytosolic intensity before and after 24 minutes LLOMe treatment in U2OS cells. Right: eGFP-2xFYVE intensity in the cytosol at different times post-induction of LMP by LLOMe. Scale bar, 10 µm. Data represent mean of 12 cytoplasmic ROIs in 12 cells ± s.e.m. See also Figure 1d. (b) Quantification of the Pearson’s Coefficient of GFP-2xFYVE and mCherry-LAMP1 from U2OS cells before and after 23 minutes treatment with control (DMSO), 0.5mM LLOMe and 0.5mM LLOMe plus 5µM VPS34-IN1. Dotted line denotes Pearson’s Coefficient of GFP-2xFYVE and mCherry-LAMP1 before treatment set to 1. One-way ANOVA, n=10 cells analysed for each condition. (c) Left: time-lapse of single lysosome in U2OS cells co-expressing GFP-OSBP-PH and mCherry-LAMP1 imaged after different durations of LLOMe treatment. Scale bar, 2 µm. Right: quantification of the eGFP-OSBP-PH intensity on lysosomes. n = 30 lysosomes quantified from 14 cells. Data are mean ± s.e.m. (d) PI(3,5)P_2_ dynamics after LLOMe treatment. Top: representative U2OS cell expressing GFP-SNXA imaged after different durations of LLOMe treatment. Scale bar, 10 µm. Bottom: the number of SNXA puncta/cell area quantified from images as shown in the top panel. Data represent mean of 3 independent experiment ± s.e.m from 33 cells. (e) Enrichment of myotubularin phosphatases in Lyso-IP fractions of 30 minutes 1mM LLOMe-treated over control (DMSO)-treated HEK293-TMEM192-3×HA cells determined by mass spectrometry. (f) Representative U2OS cells expressing GFP-MTM1, GFP-MTMR1, GFP-MTMR2, GFP-MTMR6, GFP-MTMR7, GFP-MTMR8 or YFP-MTMR14 imaged before and after 2 h LLOMe treatment. Scale bar, 10 µm. (g) Quantification of the number of MTMR14 puncta/cell from control and 1 h 1mM LLOMe treated U2OS cells. t test (n = 40 cells). (h) C2C12 cells co-expressing YFP-MTMR14 and mCherry-Galectin3 imaged after 2 h control (DMSO) or LLOMe treatment. Scale bar, 10 µm. (i) BV2 cells co-expressing YFP-MTMR14 and mCherry-Galectin3 imaged after 1 h LLOMe treatment. Scale bar, 10 µm. (j) HMC3 cells co-expressing YFP-MTMR14 and mCherry-Galectin3 imaged before and after LLOMe treatment. Scale bar, 10 µm. (k) Left: representative HeLa cells expressing YFP-MTMR14 imaged after 30 minutes treatment with different concentrations of LLOMe. Scale bar, 10 µm. Right: quantification of MTMR14 foci per cell in images as shown in the top panel. one-way ANOVA (34 cells in control, 41 cells in 100µM LLOMe, 36 cells in 250µM LLOMe, 32 cells in 500µM LLOMe, 30 cells in 1mM LLOMe). Statistical analyses were performed using GraphPad Prism. Two-tailed unpaired t-test, paired t-test or one-sample t-tests were conducted using column statistics to compare the sample means to a hypothetical value of 1. All bar graphs represent mean ± SD unless otherwise stated. ***p < 0.001, **p < 0.01, *p < 0.05.

**Extended Data Fig. 2.**
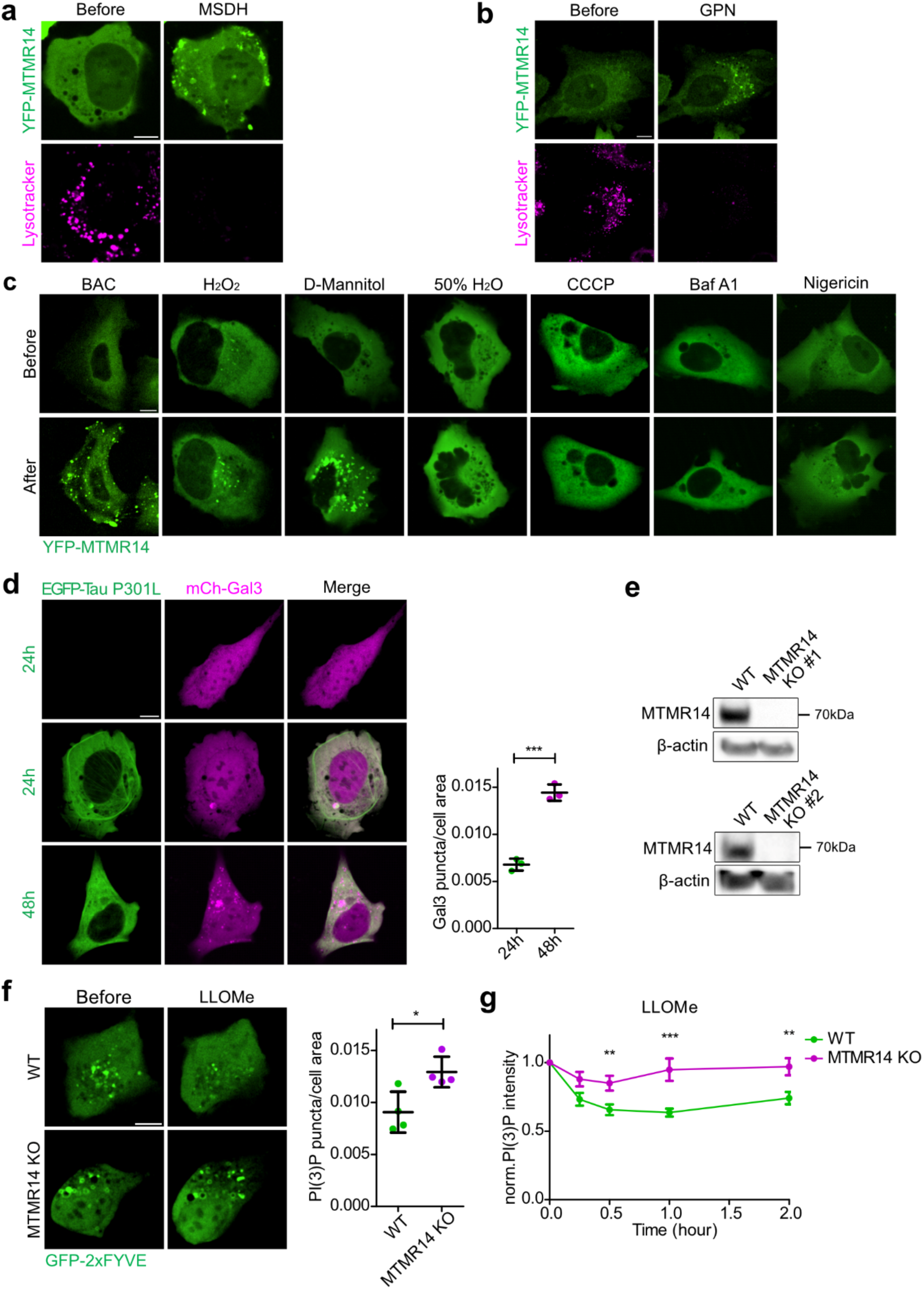
MTMR14 is recruited to cellular membranes in response to various stress conditions. (a) Representative U2OS cell expressing YFP-MTMR14 stained with LysoTracker and imaged before and after 30 minutes 50 µM MSDH treatment. Scale bar, 10 µm. (b) Representative U2OS cell expressing YFP-MTMR14 stained with LysoTracker and imaged before and after 30 minutes 200 nM GPN treatment. Scale bar, 10 µm. (c) Representative U2OS cells expressing YFP-MTMR14 imaged before and after 5 µg/mL BAC, 0.5 mM H_2_O_2_, 0.25 M D-Mannitol, 50% H_2_O, 5 µM CCCP, 100 nM Bafilomycin A1 or 5 µM Nigericin treatment. Scale bar, 10 µm. (d) Left: representative U2OS cells co-expressing GFP-Tau P301L and mCherry-Galectin3 for 24 h or 48 h. Scale bar, 10 µm. Right: quantification of Galectin3 puncta/cell area after 24 h or 48 h GFP-Tau P301L expression in U2OS cells as shown in the left panel. t test (n = 3 independent experiments, total number of cells is 86 in 24 h and 107 in 48 h conditions). (e) Top: immunoblot of U2OS WT and MTMR14 KO #1 cells. Bottom: immunoblot of U2OS WT and MTMR14 KO #2 cells. (f) Left: WT and MTMR14 KO #1 U2OS cells expressing GFP-2xFYVE imaged before and after 1 h 1mM LLOMe treatment. Scale bar, 10 µm. Right: quantification of PI(3)P puncta/cell area in WT and MTMR14 KO #1 U2OS cells as shown in the left panel. t test (n = 4 independent experiments, total number of cells is 230 for WT and 190 for MTMR14 KO). (g) Impaired LMP-induced PI(3)P dynamics in MTMR14 knockout cells. Quantification of GFP 2xFYVE intensity from confocal images of U2OS WT and U2OS MTMR14 KO cells. Cells were treated with LLOMe for the indicated time points, fixed, permeabilized, and stained with purified GFP-2xFYVE as overlay probe. One-way ANOVA (n = 5 independent experiments, each datapoint represents 15 fields of view, one field of view containing 10-20 cells, with a size of 1664 × 1664 µm). Data are mean ± s.e.m. Statistical analyses were performed using GraphPad Prism. Two-tailed unpaired t-test, paired t-test or one-sample t-tests were conducted using column statistics to compare the sample means to a hypothetical value of 1. All bar graphs represent mean ± SD unless otherwise stated. ***p < 0.001, **p < 0.01, *p < 0.05.

**Extended Data Fig. 3.**
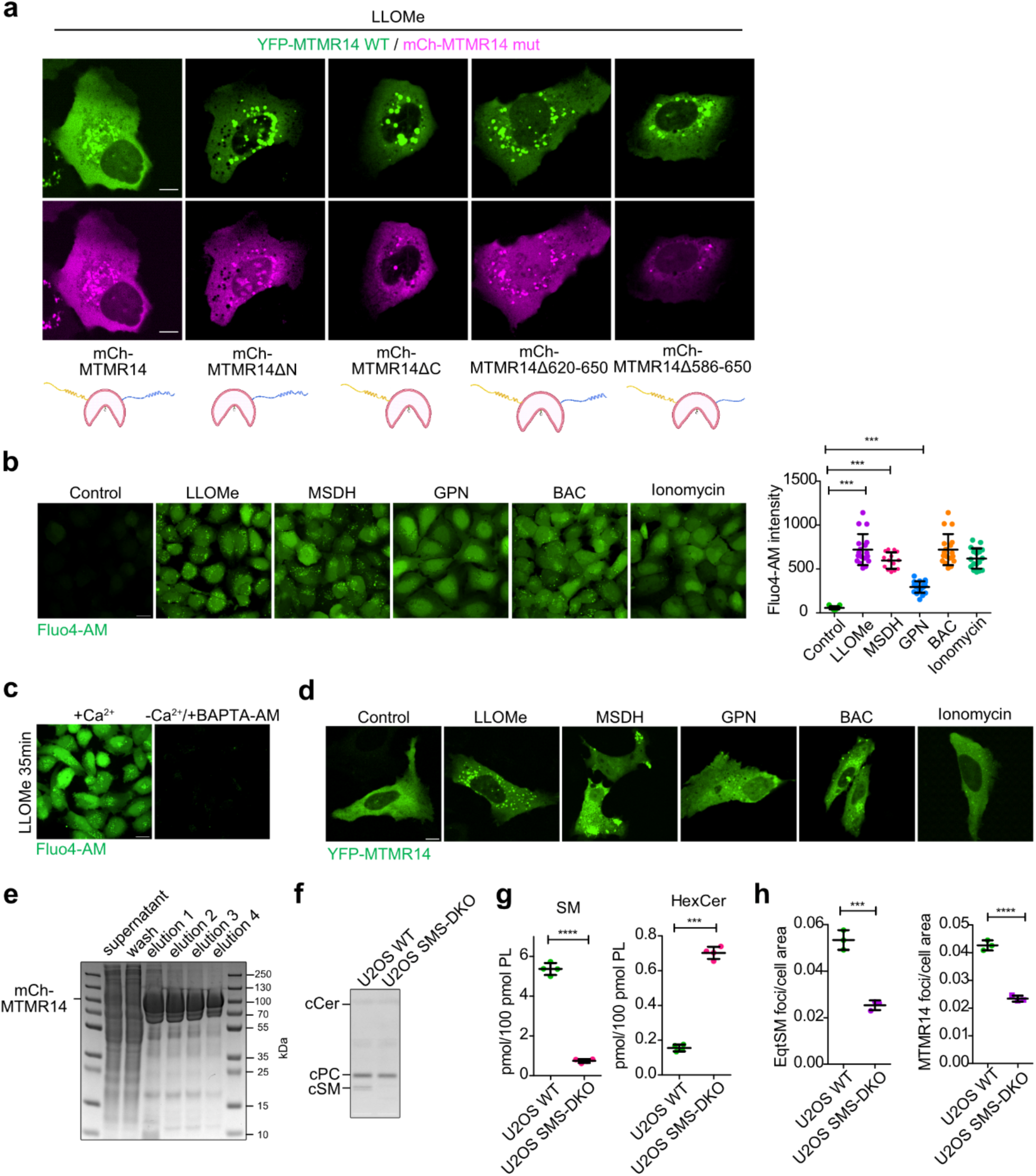
MTMR14 is recruited to damaged lysosomes by direct calcium-triggered association of a predicted alpha helix within its intrinsically disordered C-terminal domain with sphingomyelin. (a) Representative confocal images of U2OS cells co-expressing YFP-MTMR14 WT with mCherry-MTMR14 WT, mCherry-MTMR14 ΔN-term, mCherry-MTMR14 ΔC-term, mCherry-MTMR14 Δ620-650, or mCherry-MTMR14 Δ586-650 after LLOMe treatment. Scale bar, 10 µm. Lower panel: cartoons illustrating the deletion mutants with N-terminus in yellow, C-terminus in blue, and phosphatase domain in pink. (b) Left: representative confocal images of U2OS cells stained with Fluo4-AM after 40 minutes treatment with control (DMSO), LLOMe, 25 µM MSDH, 200 nM GPN, 5 µg/ml BAC, or 5 µM Ionomycin. Scale bar, 20 µm. Right: Fluo4-AM intensity quantification in images. one-way ANOVA, control (n = 17 fields of view), LLOMe (n = 18 fields of view), MSDH (n = 14 fields of view), GPN (n = 20 fields of view), BAC (n = 18 fields of view), Ionomycin (n = 17 fields of view), one field of view containing 10-20 cells, with a size of 1664 × 1664 µm. (c) Representative confocal images of U2OS cells in DMEM/FBS growth medium containing Ca^2+^ and Ca^2+^ depletion medium (2 mM EDTA in DMEM/FBS + 100 µM BAPTA-AM) stained with Fluo4-AM after 35 minutes treatment with LLOMe. Scale bar, 10 µm. (d) Representative confocal images of U2OS cells expressing YFP-MTMR14 treated with control (DMSO), LLOMe, 25 µM MSDH, 200 nM GPN, 5 µg/ml BAC, or 5 µM Ionomycin. Scale bar, 10µm. (e) Coomassie blue staining of purified mCherry-MTMR14. (f) Total lipids were extracted from WT or SMS1/2 DKO U2OS cells metabolically labeled with a clickable sphingosine analogue, click-reacted with 3-azido-7-hydroxycoumarin, separated by TLC, and analyzed by fluorescence detection. cCer, coumarin-labeled ceramide; cPC, coumarin-labeled phosphatidylcholine; cSM, coumarin-labeled sphingomyelin. (g) Levels of sphingomyelin (SM; left) and hexosylceramide (HexCer; right) in total lipid extracts from wildtype (WT) and SMS1/2 DKO cells determined by LC-MS/MS and expressed in pmol per 100 pmol of total phospholipid analyzed. t test (n = 4 independent experiments). (h) Left: number of eGFP-EqtSM puncta/cell area quantified from confocal images of U2OS WT and SMS-DKO cells after 1 h LLOMe treatment, t test (n = 3 independent experiments, total number of cells is 102 for WT and 125 for SMS-DKO). Right: number of MTMR14 puncta/cell area in confocal images of U2OS WT and SMS-DKO cells after 1 h LLOMe treatment, t test (n = 3 independent experiments, total number of cells is 117 for WT and 113 for SMS-DKO). Statistical analyses were performed using GraphPad Prism. Two-tailed unpaired t-test, paired t-test or one-sample t-tests were conducted using column statistics to compare the sample means to a hypothetical value of 1 or one-way ANOVA with Tukey’s multiple comparisons test. All bar graphs represent mean ± SD unless otherwise stated. ***p < 0.001, **p < 0.01, *p < 0.05.

**Extended Data Fig. 4.**
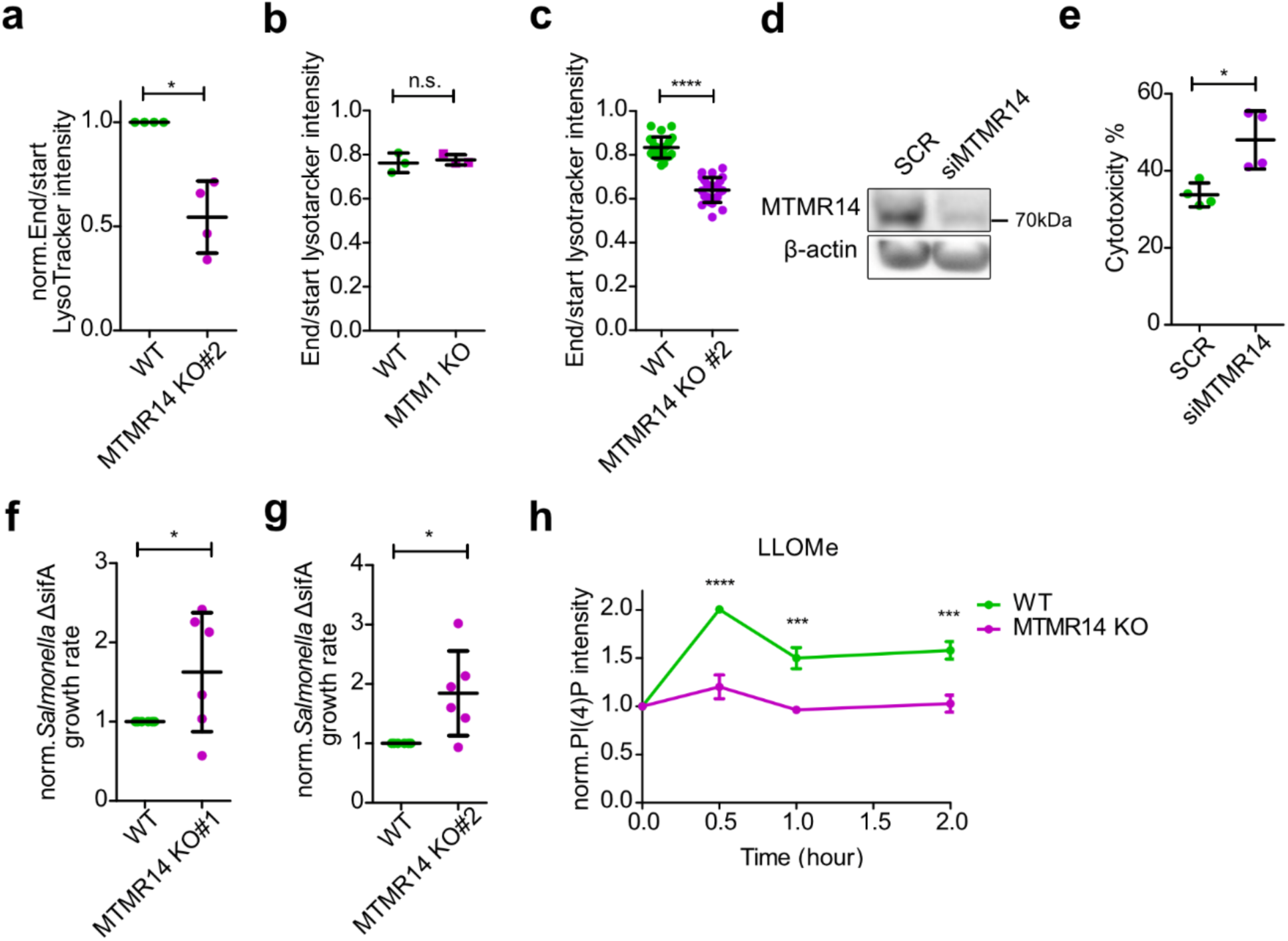
MTMR14 is required for the cellular response to LMP. (a) Quantification of mean LysoTracker intensity/field of view, fold change over control in U2OS WT and MTMR14 KO #2, t test (n = 4 independent experiments, each datapoint represents 10 fields of view, one field of view containing 35-50 cells, with a size of 3328 × 3328 µm). (b) Quantification of mean LysoTracker intensity/field of view in HeLa WT and MTM1 KO cells after 5 h chronic LLOMe treatment, t test (n = 3 independent experiments, each datapoint represents 30 fields of view, one field of view containing 10-20 cells, with a size of 1664 × 1664 µm). (c) Quantification of mean LysoTracker intensity/field of view in U2OS WT and MTMR14 KO clone #2 with 5 h chronic LLOMe treatment, t test, WT (n = 25 fields of view), MTMR14 KO (n = 25 fields of view), one field of view containing 35-50 cells, with a size of 3328 × 3328 µm. (d) MTMR14 knockdown efficiency in C2C12 cells transfected with scr control or siRNA against MTMR14. (e) Quantification of cytotoxicity using LDH assay in C2C12 cells transfected with scr control or siRNA against MTMR14, t test (n = 4 independent experiments, each datapoint represents triplicate measurements). (f) Quantification of *Salmonella* ΔsifA mutant strain growth rate (24 h post-infection) by CFU assay in U2OS WT and MTMR14 KO cells clone #1, t test. (n = 6 independent experiments). (g) Quantification of Salmonella ΔsifA mutant strain growth rate (24 h post-infection) by CFU assay in U2OS WT and MTMR14 KO cells clone #2, t test. (n = 6 independent experiments). (h) Quantification of PI(4)P intensity by immunofluorescence in U2OS WT and MTMR14 KO cells plotted as fold change over untreated. One-way ANOVA (n = 3 independent experiments, each time point in each experiment represents 20 fields of view, one field of view containing 10-20 cells, with a size of 1664 × 1664 µm). Data are mean ± s.e.m. Statistical analyses were performed using GraphPad Prism. Two-tailed unpaired t-test, paired t-test or one-sample t-tests were conducted using column statistics to compare the sample means to a hypothetical value of 1. All bar graphs represent mean ± SD unless otherwise stated. ***p < 0.001, **p < 0.01, *p < 0.05.

**Extended Data Fig. 5.**
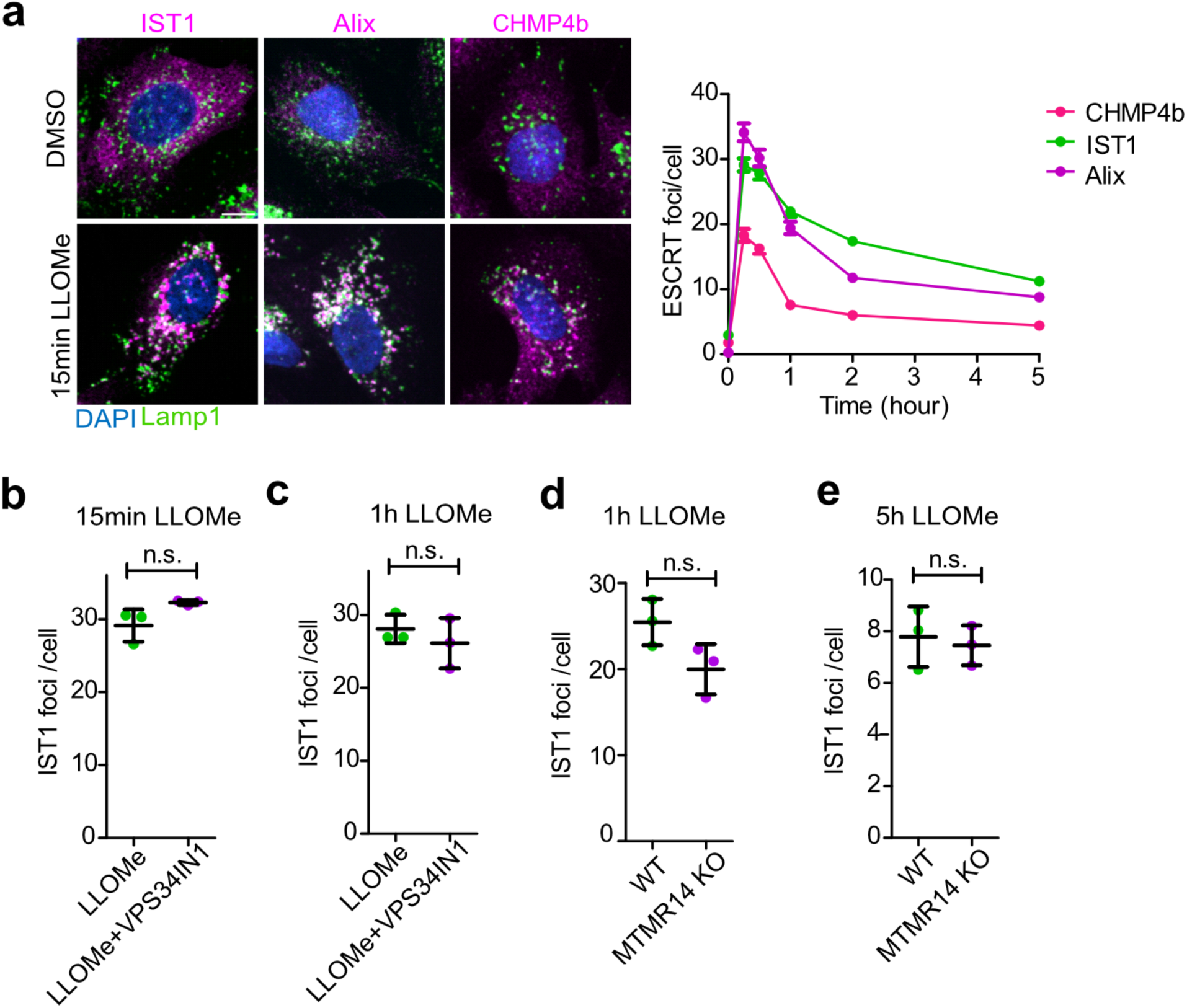
ESCRT recruitment to damaged lysosomes is independent of PI(3)P synthesis and MTMR14-mediated turnover. (a) Left: U2OS cells co-stained with DAPI, Lamp1 and either ESCRT subunit IST1, Alix, or CHMP4b after control (DMSO) or LLOMe treatment. Scale bar, 10µm. Right: quantification of ESCRT protein foci per cell upon LLOMe treatment, each datapoint represents over 200 cells. (b) Quantification of IST1 foci per cell upon 15 minutes LLOMe treatment and LLOMe plus VPS34-IN1 treatment. (n = 3 independent experiments), total number of cells is 781 in LLOMe condition, 1103 in LLOMe plus VPS34-IN1 condition. (c) Quantification of IST1 foci per cell upon 1 h LLOMe treatment and LLOMe plus VPS34-IN1 treatment. (n = 3 independent experiments), total number of cells is 1076 in LLOMe condition, 1020 in LLOMe and VPS34-IN1 condition. (d) Quantification of IST1 foci in U2OS WT and MTMR14 KO cells upon 1h LLOMe treatment, t test (n = 3 independent experiments), total number of cells is 535 in WT, 471 in KO cells. (e) Quantification of IST1 foci in U2OS WT and MTMR14 KO cells upon 5h LLOMe treatment, t test (n = 3 independent experiments), total number of cells is 387 in WT, 411 in KO cells. Statistical analyses were performed using GraphPad Prism. Two-tailed unpaired t-test, paired t-test or one-sample t-tests were conducted using column statistics to compare the sample means to a hypothetical value of 1. All bar graphs represent mean ± SD unless otherwise stated. ***p < 0.001, **p < 0.01, *p < 0.05.

**Extended Data Fig. 6.**
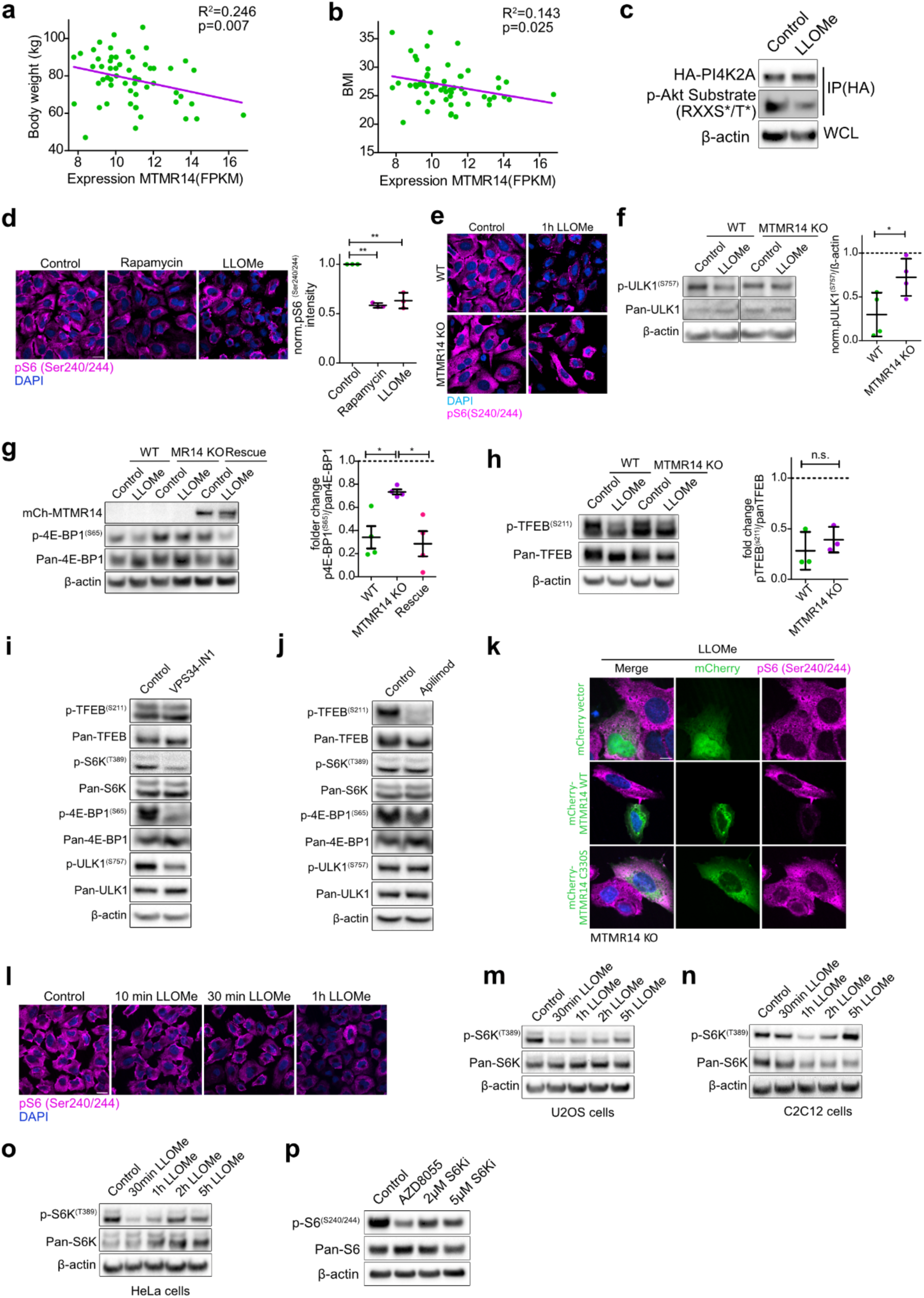
Inhibition of mTORC1-S6K signaling rescues defective lysosome repair in MTMR14 KO cells. (a) MTMR14 expression in human skeletal muscle inversely correlates with body weight. FPKM: fragments per kilobase per million mapped fragments, data were collected from 56 samples. (b) MTMR14 expression in human skeletal muscle. Linear plots representing the correlation of MTMR14 expression in human skeletal muscle samples and body mass index (BMI). FPKM: fragments per kilobase per million mapped fragments, data come from 56 samples. (c) LLOMe treatment decreases PI4K2A phosphorylation. Immunoprecipitation (IP) fractions of U2OS cells expressing HA-PI4K2A and treated with DMSO control or 30 minutes LLOMe were collected with anti-HA beads and immunoblotted with antibodies against the p-RXRXXs/t* motif. (d) Left: representative confocal images of U2OS cells treated with control (DMSO), 1 h 1 µM rapamycin, or 1 h 1 mM LLOMe stained with DAPI and antibodies against pS6 (Ser240/244). Scale bar, 20µM. Right: quantification of pS6 (Ser240/244) intensity. t test (n = 3 independent experiments, each datapoint represents 15 fields of view, one field of view containing 10-20 cells, with a size of 1664 × 1664 µm). (e) Representative confocal images of U2OS and MTMR14 KO cells treated with control (DMSO) or 1 h 1 mM LLOMe stained with DAPI and antibodies against pS6 (Ser240/244). Scale bar, 20µM. Quantification is shown in Figure 4c. (f) Left: Representative immunoblot of control and LLOMe treated WT and MTMR14 KO U2OS cell lysates. Right: quantitative analysis of pULK1 S757/ total ULK1 levels in control or LLOMe-treated (2 h) WT and MTMR14 KO U2OS cells. t test (n = 4 independent experiments). Dotted line denotes pULK1 S757/ total ULK1 levels in DMSO controls set to 1. (g) MTMR14 re-expression in KO cells rescues decreased 4E-BP1 phosphorylation upon LLOMe treatment. Left: representative immunoblot of U2OS WT, MTMR14 KO, and MTMR14 KO cells expressing mCherry-MTMR14 under doxycycline control subjected to 2 h DMSO control or LLOMe treatment. Right: densitometric quantification of 4E-BP1 S65 phosphorylation relative to total 4E-BP1. one-way ANOVA (n = 4 independent experiments). Dotted line denotes p4E-BP1 S65/ total 4E-BP1 in controls set to 1. (h) Left: representative immunoblot of U2OS WT and MTMR14 KO cells treated for 2 h with DMSO control or LLOMe. Right: quantification of pTFEB S211 relative to total TFEB from immunoblots as shown on the left. t test (n = 3 independent experiments). Dotted line denotes pTFEB S211/ total TFEB in controls set to 1. (i) Immunoblot of U2OS cells treated for 1 hr with DMSO control or 5 µM VPS34-IN1. (j) Immunoblot of U2OS cells treated for 1 hr with DMSO control or 200 nM Apilimod. (k) Representative confocal images of 1 h 1 mM LLOMe treated U2OS MTMR14 KO cells expressing mCherry-vector, mCherry-MTMR14 WT, or mCherry-MTMR14 C330S stained with DAPI and antibodies against pS6 (Ser 240/244). Scale bar, 10 µm. Quantification is shown in Figure 4e. (l) Representative confocal images of DMSO control or different time points of 1 mM LLOMe treated U2OS stained with DAPI and antibodies against pS6 (Ser 240/244). Quantification is shown in Figure 4f. (m) Immunoblot of U2OS cells treated with DMSO control or different time points of LLOMe. (n) Immunoblot of C2C12 cells treated with DMSO control or different time points of LLOMe. (o) Immunoblot of HeLa cells treated with DMSO control or different time points of LLOMe. (p) AZD8055 and S6Ki efficiency test. Immunoblot of S6 S240/S244 phosphorylation in U2OS cells treated for 1 h with DMSO control, 200 nM AZD8055, 2 µM S6Ki, or 5 µM S6Ki. Statistical analyses were performed using GraphPad Prism. Two-tailed unpaired t-test, paired t-test or one-sample t-tests were conducted using column statistics to compare the sample means to a hypothetical value of 1. All bar graphs represent mean ± SD unless otherwise stated. ***p < 0.001, **p < 0.01, *p < 0.05.

**Extended Data Fig. 7.**
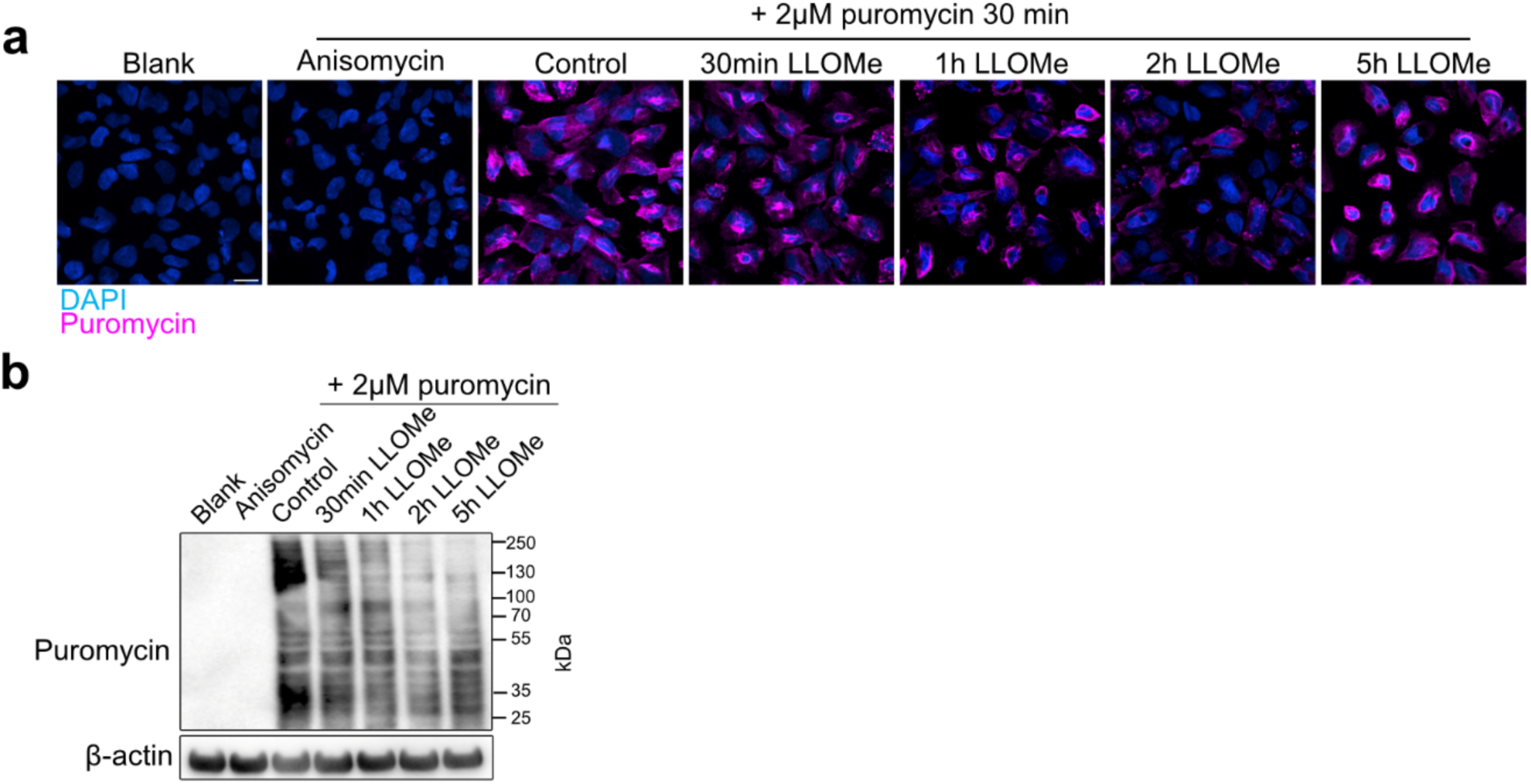
Time-dependent inhibition of protein synthesis following lysosomal damage. (a) Representative confocal images of LLOMe-treated U2OS cells stained with 2µM puromycin and DAPI. Quantification is shown in Figure 5a. (b) Immunoblot of U2OS cells treated with DMSO control or different time points of LLOMe. Statistical analyses were performed using GraphPad Prism. Two-tailed unpaired t-test, paired t-test or one-sample t-tests were conducted using column statistics to compare the sample means to a hypothetical value of 1. All bar graphs represent mean ± SD unless otherwise stated. ***p < 0.001, **p < 0.01, *p < 0.05.

**Extended Data Fig. 8.**
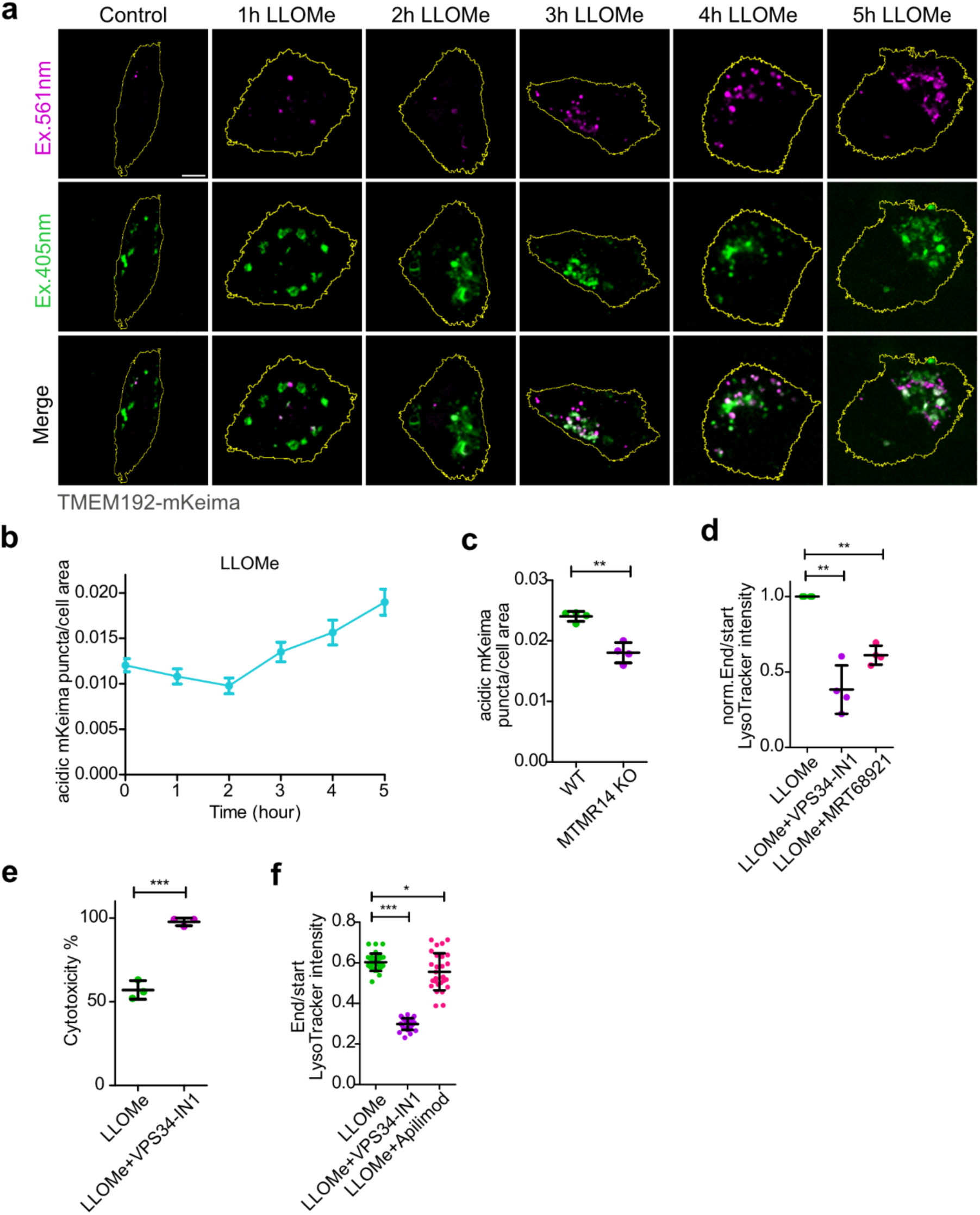
PI(3)P synthesis and turnover control ULK1-dependent lysophagy. (a) Representative confocal live cell images of TMEM192-mKeima expressing HeLa cells treated with LLOMe for the indicated time points. Quantification is shown in Figure 6a. (b) Quantification of acidic TMEM192-mKeima puncta/cell area in U2OS cells after different durations of LLOMe treatment. n = 28-37 cells per datapoint. Data are mean ± s.e.m. (c) Quantification of acidic TMEM192-mKeima puncta/cell area after 5 h LLOMe treatment in U2OS WT and MTMR14 KO cells. t test (n = 4 independent experiments, total number of cells is 116 for WT and 113 for MTMR14 KO). Representative confocal images in Figure 6e. (d) Quantification of mean LysoTracker intensity/field of view fold change over control in BV2 cells after 5 h LLOMe plus control (DMSO), 5 µM VPS34-IN1, or 2µM MRT68921 treatment. one-way ANOVA (total number of fields of view is 30 each group, one field of view containing 35-50 cells, with a size of 3328 × 3328 µm). (e) Quantification of LDH cytotoxicity assay in BV2 cells treated for 5 h with 4mM LLOMe plus DMSO control or 5 µM VPS34-IN1. t test (n = 3 independent experiments, each datapoint represents triplicate measurements). (f) Quantification of mean LysoTracker intensity/field of view fold change over control in HeLa cells after 5 h LLOMe plus control (DMSO), 5 µM VPS34-IN1, or 200 nM Apilimod treatment. one-way ANOVA (total number of fields of view is 30 for LLOMe, 29 for LLOMe + VPS34-IN1, and 28 for LLOMe + Apilimod, one field of view containing 35-50 cells, with a size of 3328 × 3328 µm). Statistical analyses were performed using GraphPad Prism. Two-tailed unpaired t-test, paired t-test or one-sample t-tests were conducted using column statistics to compare the sample means to a hypothetical value of 1 or one-way ANOVA with Tukey’s multiple comparisons test. All bar graphs represent mean ± SD unless otherwise stated. ***p < 0.001, **p < 0.01, *p < 0.05.

**Extended Data Fig. 9.**
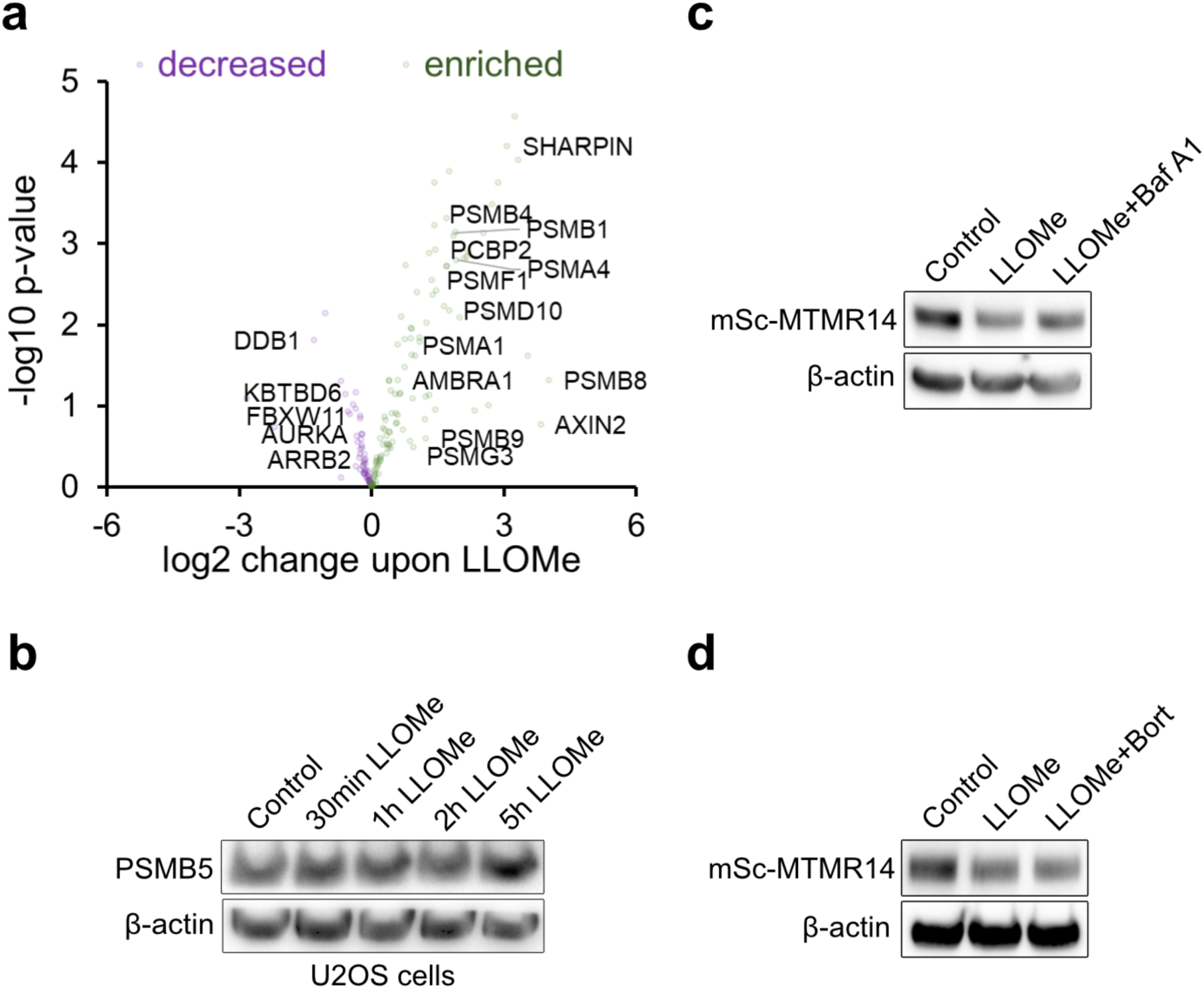
Proteasome recruitment and ubiquitination in response to lysosomal damage. (a) Volcano plot illustrating the enrichment of proteasome subunits and ubiquitylation machinery on lysosomes immunoprecipitated from control or 30min 1mM LLOMe-treated HEK293-TMEM192-3xHA cells determined by quantitative proteomics. (b) Immunoblot of U2OS cells treated with DMSO control, or 1mM LLOMe for 30 minutes, 1 h, 2 h, or 5 h. (c) Representative immunoblot of U2OS-mScarlet-MTMR14 KI cells treated with DMSO control, 5 h 1mM LLOMe, and 5 h LLOMe with 200 nM Baf A1. (d) Representative immunoblot of U2OS-mScarlet-MTMR14 KI cells treated with DMSO control, 5 h 1mM LLOMe, and 5 h LLOMe with 2 µM Bortezomib. Statistical analyses were performed using GraphPad Prism. Two-tailed unpaired t-test, paired t-test or one-sample t-tests were conducted using column statistics to compare the sample means to a hypothetical value of 1 or one-way ANOVA with Tukey’s multiple comparisons test. All bar graphs represent mean ± SD unless otherwise stated. ***p < 0.001, **p < 0.01, *p < 0.05.

## Legends to supplementary videos

**Video S1.** Live imaging of MTMR14 recruitment to damaged lysosomes in HeLa cells following LLOMe treatment.

Live-cell spinning disk confocal imaging of HeLa cells co-expressing YFP-MTMR14 and mCherry-Gal3. Videos were acquired at 1 frame/ 10 min for 120 min.

**Video S2.** Live imaging of LLOMe-induced recruitment kinetics of mCherry-MTMR14 C-terminus and PI(4)P sensor GFP-OSBP-PH co-expressed in HeLa cells. Videos were acquired at 1 frame/ 5 min for 40 min.

## Legends to supplementary tables

### Supplementary Table 1

Lyso-IP MS/MS analysis of lysosome-IP samples from HEK293 cells treated with LLOMe (1 mM) for 30 min.

List of all the identified proteins in HEK293 cells stably expressing TMEM192-3×HA comparing DMSO and LLOMe (30 min, 1mM). Related to Figures 1E,; Figure S1I, S8O. The table comprises log10 intensity as well as the normalized mean and p values.

### Supplementary Table 2

Lyso-IP MS/MS analysis of lysosome-IP samples from HEK293 cells treated with LLOMe (1 mM) for 1h.

List of identified proteins in HEK293 cells stably expressing TMEM192-3×HA compared DMSO and LLOMe (1h, 1mM). Related to Figure 7A. The table comprises log10 intensity as well as the normalized mean and p values.

## Notes

### Competing Interest Statement

The authors have declared no competing interest.

